# A human immune system mouse model for preclinical evaluation of therapies in pemphigoid disease

**DOI:** 10.64898/2026.01.15.699625

**Authors:** Leonie Voss, Katharina Maier, Stella Wagner, Ann-Kathrin Schneider, Mareile Schlotfeldt, Leon Altmann, Emir Ucgan, Lukas Hönninger, Matthias Peipp, Falk Nimmerjahn, Katja Bieber, Anja Lux

**Affiliations:** Division of Genetics, Department of Biology, Friedrich-Alexander-Universität Erlangen-Nürnberg; Erlangen, Germany; Profile Center for Immunomedicine (I-MED), Friedrich-Alexander-Universität Erlangen-Nürnberg; Erlangen, Germany; Stem Cell Transplantation and Immunotherapy, Division of Antibody-Based Immunotherapy, Department of Medicine II, Christian Albrechts University Kiel and University Medical Center Schleswig-Holstein; Kiel, Germany; Lübeck Institute of Experimental Dermatology, University of Lübeck; Lübeck, Germany

## Abstract

Pemphigoid diseases (PD) including Epidermolysis bullosa acquisita (EBA) are rare immunoglobulin G (IgG)-driven autoimmune skin blistering diseases with limited therapeutic options. Mechanistically, chronic inflammation in the skin leads to disruption of the dermal-epidermal junction (DEJ) with a crucial contribution of FcγR-mediated activation of myeloid immune cells such as neutrophils. Thus, targeting of kinases involved in FcγR-dependent activation of myeloid immune cells holds great promise as a therapeutic strategy. In this study, we employ human immune system (HIS) mice featuring all major human leukocytes which allows to investigate the impact of therapeutics on human immune cells in vivo. In a passive transfer approach, repetitive application of collagen VII (COL7^c^) specific IgG was associated with skin inflammation including infiltration of activated human immune cells and thickening of the epidermis. While application of recombinant human G-CSF boosted myeloid cell maturation and thus disease severity, treatment with FcγR-blocking antibodies impaired disease development confirming the crucial role of human cells. Finally, small molecule PDK1 inhibitor BX-795 abrogated development of skin inflammation associated with reduced leukocyte infiltration and activation supporting the role of PDK1 in FcγR-driven immune cell activation. This study establishes the first in vivo model of EBA in HIS mice and reveals its suitability for pre-clinical screening and evaluation of therapeutic agents. Importantly, it highlights the potential of kinase inhibition for treatment of EBA.

## INTRODUCTION

Pemphigoid diseases (PD) such as epidermolysis bullosa acquisita (EBA) are a rare form of autoimmune disease caused by autoantibodies against components of the dermal-epidermal junction (DEJ). In EBA, collagen VII (COL7^c^) specific immunoglobulin G (IgG) antibodies bind to the DEJ and lead to inflammation via recruitment of immune cells involving both IgG Fc receptors as well as the complement system (*1–4*).

Mechanistically, deposited IgG acts as an immune complex (IC) able to crosslink FcγRs on myeloid effector cells (*5*). These FcγRs can then trigger either activating or inhibitory intracellular signaling cascades. Activating FcγRs in humans are FcγRI (CD64), FcγRIIa (CD32A) and FcγRIIIa (CD16A). While FcγRIIa itself carries the immunoreceptor tyrosine-based activation motif (ITAM) required for signal transduction, FcγRI and FcγRIIIa depend on association with the signaling accessory FcRγ chain for their function (*6*). Inhibitory signaling is induced by FcγRIIb (CD32B) which includes an immunoreceptor based inhibitory motif (ITIM). In addition, human immune cells express FcγRIIIb (CD16B) which lacks a signaling motif altogether due to its glycosyl-phosphatidylinositol (GPI) anchor. Most importantly, immune cells regularly co-express several different FcγRs allowing fine-tuning of effector responses depending on receptor expression levels and engagement by IgG. Notable exceptions include B cells, which solely express FcγRIIb and T cells which lack FcγR expression (*7*). Signaling cascades triggered downstream of activated FcγRs prominently involve kinases from the Src and the Spleen tyrosine kinase (Syk) family. The Src family kinases serve as essential initiators of receptor activation by phosphorylation of ITAMs leading to recruitment and activation of Syk kinases. Ultimately, these signaling cascades are crucial for immune cell activation upon IgG crosslinking as they induce a multitude of effector functions driving inflammatory responses. This includes release of inflammatory mediators, uptake of the immune complex by phagocytosis potentially followed by processing and presentation of the antigen or secretion of cytotoxic agents in a cell-type specific manner (*8*). While beneficial in protective immune responses, these effector functions cause destruction of tissue and chronic inflammation when directed against self-antigens as is the case in autoimmunity. With respect to EBA, production of reactive oxygen species (ROS) appears to be particularly important as their secretion will directly affect the DEJ causing its dissolution and formation of skin blisters (*5, 9*).

Current treatment of PD/EBA is challenging and typically involves a combination of anti-inflammatory, immunosuppressive and immunomodulating agents (*10*). Systemic corticosteroids like Prednisolone are often used as first-line treatment. If steroids alone are insufficient, anti-inflammatory agents like Dapsone or immunosuppressants such as Azathrioprine are added (*11–13*). Intravenous immunoglobulin G (IVIG) described to have anti-inflammatory properties upon high-dose treatment, has also been applied in EBA therapy when the disease is severe and refractory to conventional treatments (*14–16*). As current treatments are rather broad, have significant side effects and do not consistently lead to long-term remission, further therapeutic targets are urgently needed. The prominent role of myeloid effector cells responding to IgG autoantibodies indicates that inhibition of the kinases involved in FcγR signaling might be a promising strategy in inflammatory disorders of the skin. Such small signal transduction inhibitors e.g. targeting PI-3-Kinase (PI3K), Brutońs tyrosine kinase (BTK), Syk or janus kinases (JAK) are already licensed for therapy of various cancers and hold significant potential for managing chronic inflammatory diseases (*17–22*). 3-phosphoinositide-dependent kinase 1 (PDK1) is a key signaling molecule within the PI3K pathway located upstream of Akt (*23*) and has been described to maintain immune cell development and function (*24*). Importantly, PI3K is induced by FcγRs in myeloid cells (*25*) and PDK1/Akt signaling was shown to be critical for several neutrophil functions, including chemotaxis, degranulation and ROS production highlighting its potential as a therapeutic target in EBA (*26, 27*).

Research on pathogenesis of PD and pre-clinical assessment of therapeutic agents in the context of human immunity is severely impaired by a lack of in vivo model systems reflecting the genetic complexity of the human immune system. Currently, the pathogenic potential of human IgG autoantibodies and/or patient-derived leukocytes as well as therapeutic interventions are often investigated ex vivo applying human skin organ cultures or cryosection assays (*5, 16, 28*). In vivo pathogenicity of human IgG from EBA patients could be demonstrated upon passive transfer into mice but patient-derived sera are rare and results are heterogenous indicating limited cross-reactivity (*29, 30*). Most importantly, disease development in established in vivo models is mediated by murine effectors, underscoring the need for alternative animal models that allow to dissect human-specific mechanisms. As recombinant pathogenic COL7^c^-specific IgG are lacking, the most widely used model for EBA induction is the passive transfer of rabbit-derived mouse COL7^c^ specific IgG (*31, 32*). This approach allows to mimic the effector phase of the disease and led to discoveries on the role of FcγR (*33*), neutrophils (*5*) but also Th17 cells (*32*) in EBA development. While undoubtedly crucial, such in vivo studies in mice must be considered carefully as pronounced differences in FcγR expression and immune cell composition exist between mice and humans. Most importantly, human but not mouse neutrophils express FcγRIIIb and expression of inhibitory FcγRIIb on human monocytes and neutrophils is lower than on their murine counterparts. Moreover, polymorphic variants exist for human FcγRs that have been associated with altered IgG binding and/or cell activation (*7, 34, 35*). Thus, investigating efficacy and potential side effects of immunotherapeutics for IgG-driven diseases would benefit from models as close to the human immune system as possible.

We thus aimed to develop an in vivo model of EBA in which disease is mediated by human FcγRs expressed on human immune cells. Human immune system (HIS) mice are generated by transplantation of severely immunocompromised mice with human hematopoietic stem cells (HSC) and are ideally suited because of long-term presence of all major human leukocyte populations in blood and lymphoid as well as non-lymphoid organs (*7*). Induction of adaptive immune responses in HIS mice is limited which prevents active development of autoimmunity. Autoantibodies can, however, be passively administered allowing to investigate the effector phase of inflammation including clinical efficacy of drugs targeting the involved effector functions of myeloid cells (*36, 37*).

In the present study, we established a HIS model of EBA, evaluated contribution of human immune cells in disease development and assessed the potential of PDK1 inhibition for treatment.

## RESULTS

### Rabbit collagen VII-specific IgG binds to and activates human FcγRs

To allow induction of EBA in HIS mice, we used affinity-purified polyclonal antibodies from serum of immunized rabbits mimicking the standard experimental model of passive EBA in mice. While rabbit IgG had previously been shown to bind to human FcγRs (*38–40*), we aimed to first confirm efficient binding to human leukocytes and, more importantly, their subsequent activation. Applying established CHO cell lines stably expressing human FcγRs (*41, 42*) we successfully demonstrated binding of monomeric rabbit IgG to high-affinity FcγRI, -131Arg and -131His variants of FcγRIIa, -158F and -158V variants of FcγRIIIa and -NA1 and -NA2 variants of FcγRIIIb. In contrast, no significant binding to inhibitory FcγRIIb could be observed which is in line with previous reports (*40*) (Fig. 1A). Accordingly, binding of rabbit IgG to human neutrophils, monocyte subsets and NK cells could be observed while binding to B cells, and T cells was minimal (Fig. 1B). Beyond binding we further determined immune cell activation, specifically production of ROS and secretion of cytokines as these effector mechanisms are involved in EBA disease pathogenesis (*10*). Upon stimulation of cells with immobilized, plate-bound anti-COL7^c^ IgG, intracellular staining of ROS confirmed significant activation of neutrophils, classical monocytes and NK cells (Fig. 1C) while elevated concentrations of key inflammatory mediators IL-1β, IL-18 and IL-8 were identified in the cell culture supernatant (Fig. 1D).

**Fig. 1.**
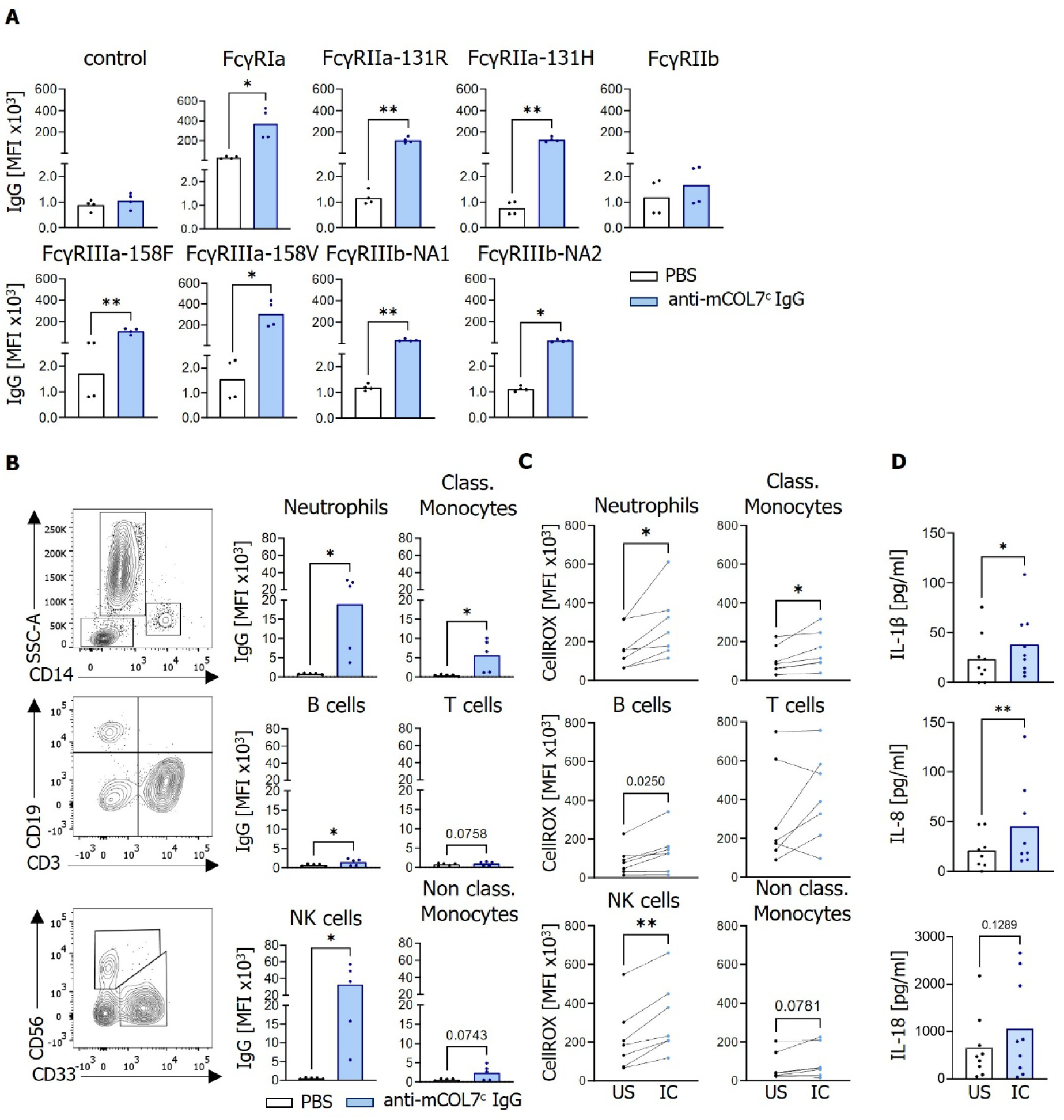
Anti-mCOL7^c^ IgG binding to human FcγRs and activation of human cells by COL7^c^ immune complexes. **(A)** Anti-mCOL7^c^ IgG binding to CHO cells stably transfected with human FcγRs and their indicated polymorphic variants. Bound IgG was detected by staining with fluorescently labelled anti-rabbit IgG and median fluorescence intensity (MFI) is depicted for n=4 independent experiments. **(B)** Flow cytometry gating and binding of anti-mCOL7^c^ IgG to human peripheral blood cells. Class., classical. Cells were pre-gated for all leukocytes (CD45+), single cells, and live cells followed by distinction of neutrophils (SSC^high^) and classical (class.) monocytes (CD14^+^). Within SSC^low^/CD14^-^ cells, B cells (CD19^+^) and T cells (CD3^+^) were selected and CD19^-^CD3^-^cells were further divided into NK cells (CD56^+^) and non-classical monocytes (CD33^+^). **(C)** Flow cytometry quantification of cytosolic ROS production for indicated human peripheral blood leukocyte populations following stimulation with plate-bound COL7^c^ immune complexes for 30min at 37°C. US, unstimulated, IC, immune complex. Cells were pre-gated for all leukocytes, single cells, and live cells. **(D)** Quantitative analysis of secreted cytokines in supernatants from immobilized COL7^c^-IC stimulated human cells, measured with the LEGENDplex^TM^ Human inflammation kit 1. Symbols indicate individual measurements or mice; bars represent the mean. Statistical significance was assessed using Shapiro-Wilk normality test followed by paired t-tests (gaussian distribution) or Wilcoxon tests (non-gaussian distribution). *p < 0.05; **p < 0.01.

### Rabbit COL7^c^-specific IgG induces skin inflammation in HIS mice

We next developed the experimental setup for induction of EBA in human immune system (HIS) mice. As described before, severely immunocompromised mice can be reconstituted with a human immune system upon transfer of human hematopoietic stem cells (HSC) (*43–45*). In this study we employed NSG/FcRγ^-/-^ mice lacking murine B cells, T cells and NK cells due to their genetic background and deletion of *il2rg* leading to impairment of cytokine receptor signaling (*46*). In addition, this mouse strain carries a deletion of *fcerg1* which encodes for the accessory chain of FcγRs required for their surface expression and signal transduction (*6*) thus abrogating IgG activity via murine activating FcγRs (Suppl. Fig. S3). Irradiation and transplantation of newborn mice with human HSCs resulted in development of human leukocyte populations within 3 to 4 months with an average reconstitution efficiency of 25% (Suppl. Fig. S1, S2 and Table 1). Disease was induced by repetitive i.p. injections of rabbit anti-COL7^c^ IgG and clinical scoring performed to monitor development of skin inflammation for 13 days (Fig. 2A). Initial experiments indeed revealed enhanced formation of scuffing and skin reddening in HIS mice compared to control mice not reconstituted with HSCs (Suppl. Fig. S3). However, key features of EBA such as separation of dermis and epidermis and thus skin blistering could not be observed. We hypothesized that low disease activity was caused by insufficient presence or maturity of human myeloid effector cells, particularly neutrophils. We thus added treatment with recombinant pegylated human G-CSF previously described to boost neutrophil maturation and egress from the bone marrow in HIS mice (*47, 48*) and observed elevated blood neutrophils but also classical monocytes (Suppl. Fig. S4). Indeed, this led to more pronounced disease as seen by an increased proportion of affected skin (Fig. 2B) upon comparable deposition of COL7^c^-specific IgG at the DEJ (Suppl. Fig. S5). While separation at the DEJ could now be identified occasionally, skin lesions were still rare suggesting that further improvements might be required for closer resemblance to the human disease. Treatment with rabbit anti-COL7^c^ IgG but not control IgG was associated with influx of cells into the skin and increased epidermal thickness which was further enhanced by co-treatment with G-CSF in line with disease severity (Fig. 2C). Flow cytometry analysis of affected skin identified infiltrating cells as classical monocytes and neutrophils with neutrophil numbers directly correlated with disease score. Infiltrating cells were also found to express CD69 (Fig. 2D). Serving as an early activation receptor, CD69 can modulate inflammatory and regulatory responses like cytokine production, cell signaling and effector functions (*49–51*) as well as contribute to myeloid cell recruitment and activation (*52*). Of note, no significant infiltration of murine leukocytes could be detected in line with their lack of expression of activating FcγRs (Fig. 2E). To investigate systemic inflammation, human cytokines were quantified in plasma. Partially independent of G-CSF addition, injection of anti-COL7^c^ IgG led to increased concentrations of multiple proinflammatory cytokines during disease development (Fig. 2F, Suppl. Fig. S6) while no significant alterations in murine cytokines could be observed (Suppl. Fig. S7).

**Fig. 2.**
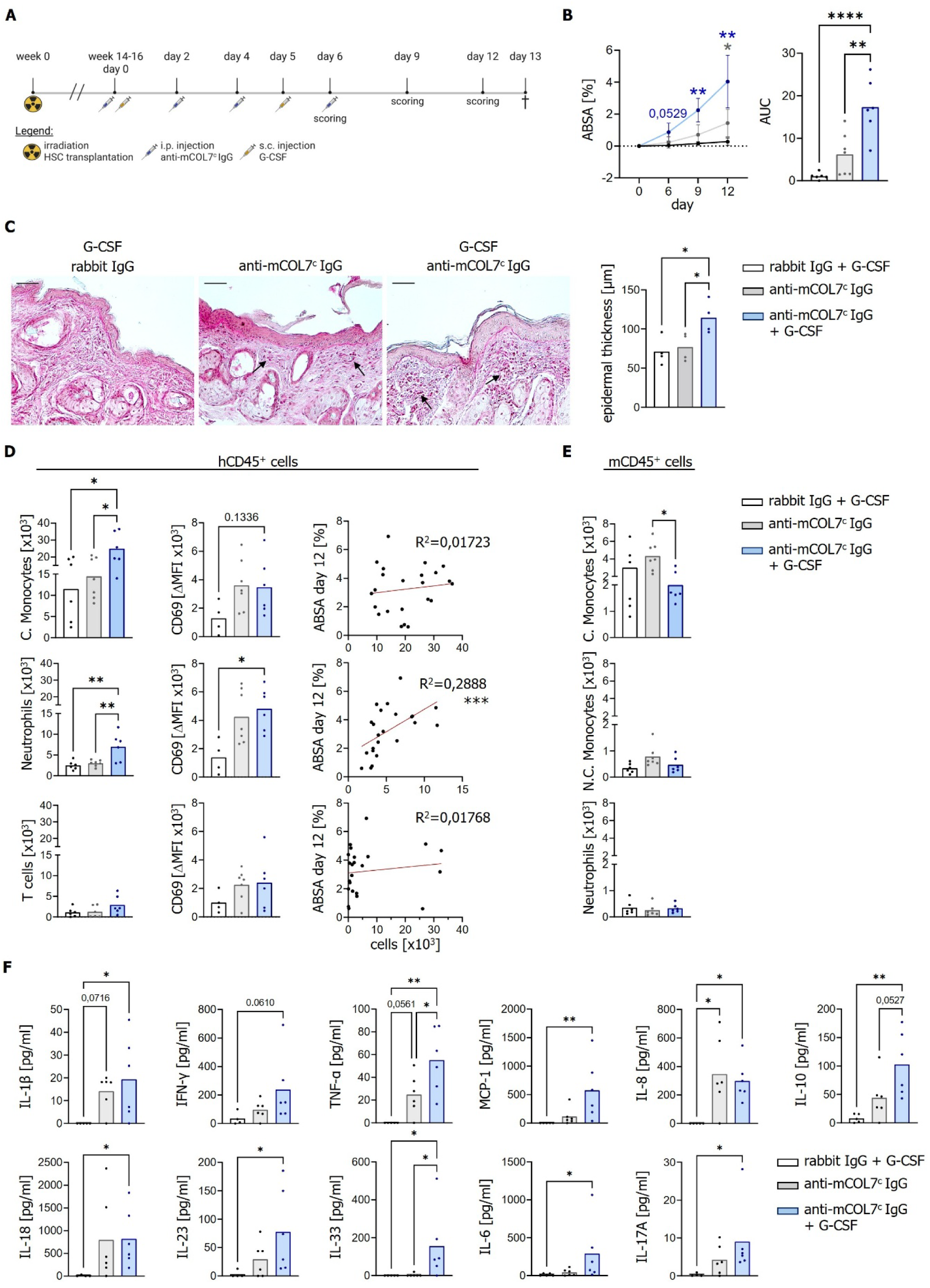
Characterization of EBA-like autoimmunity in human immune system mice. (**A**) Schematic representation of EBA induction in human immune system (HIS) mice by multiple intraperitoneal (i.p.) injections of rabbit anti-mCOL7^c^ IgG. Additional subcutaneous (s.c.) administration of recombinant human G-CSF is indicated. (**B**) EBA induction in humanized NSG/FcRγ^-/-^ mice compared to controls injected with non-specific rabbit IgG. Skin disease severity is shown as affected body surface area (ABSA, %) over time (left) and as area under the curve (AUC, right). (**C**) Exemplary images of the histological examination of skin sections using PAS staining to assess tissue structure and infiltration, marked by arrows. Epidermal thickness was quantified. (**D**) Flow cytometric analysis of human immune cell subsets (per gram skin and 100,000 live cells) and activation marker CD69 expression in skin and correlation analysis of cell counts and ABSA score on day 12. ΔMFI, median fluorescence intensity after subtraction of FMO control. (**E**) Quantification of murine immune cell populations in skin by flow cytometry (per gram skin and 100,000 live cells). (**F**) Quantification of human cytokines in plasma on day 13 after disease induction using the LEGENDplex^TM^ Human Inflammation Panel 1. Symbols indicate individual mice; bars represent the mean. Statistical analyses were performed using Shapiro-Wilk normality test followed by Ordinary one-way ANOVA (gaussian distribution) or Kruskal-Wallis test (non-gaussian distribution). *p < 0.05; **p < 0.01; ***p < 0.001; ****p <0.0001.

### Disease development requires human FcγRs

Human immune system mice are chimeras with mixed populations of human and murine immune cells. As murine leukocytes were not found to significantly infiltrate the skin during disease, we wanted to confirm contribution of human cells and, more specifically, human FcγRs. We therefore performed disease induction in presence of FcγR-specific antibodies 10.1, IV.3 and 3G8 blocking human FcγRI, FcγRIIa/b and FcγRIIIa/b, respectively (Fig. 3A) (*53–55*). This additional treatment significantly impaired skin inflammation (Fig. 3B), reduced infiltration of cells into the skin and epidermal thickness (Fig. 3C). Both the number of infiltrating classical monocytes and neutrophils as well as their activation based on expression of CD69 and CD18, a β2-integrin essential for recruitment and adhesion of myeloid effector cells to ICAM-1-expressing endothelial cells and keratinocytes (*56, 57*), were reduced (Fig. 3D). In contrast, numbers of murine immune cells in the skin were unaffected (Fig. 3E). In terms of systemic inflammation, plasma concentration of inflammatory TNFα and MCP-1, amongst others, were decreased while e.g. IFNγ were largely unaffected (Fig. 3F, Suppl. Fig. S6 and S7) which could indicate contribution of additional effector mechanisms.

**Fig. 3.**
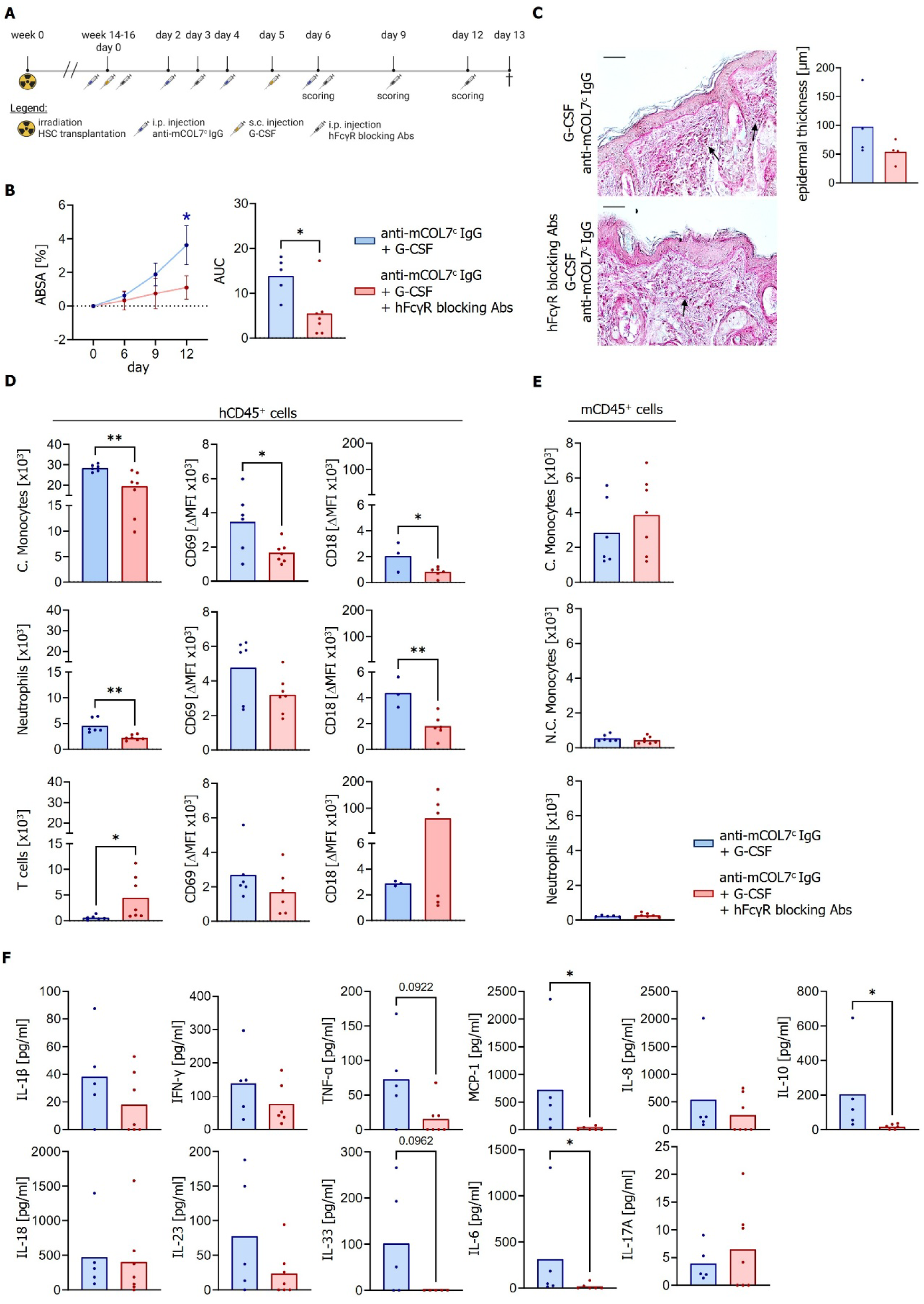
Blocking of human Fcγ receptors during induction of EBA-like autoimmunity impairs disease development in human immune system mice. (**A**) Schematic representation of the experimental approach for inducing an EBA-like phenotype in HIS mice, combined with intraperitoneal (i.p.) administration of monoclonal antibodies (clones 10.1, AT10, 3G8) to block human FcγRs. (**B**) Assessment of skin disease severity shown as % ABSA over time (left) and AUC (right). (**C**) Histological analysis of skin sections using PAS staining for tissue structure and infiltration (arrows) evaluation (left) and quantification of epidermal thickness (in µm, right). (**D**) Flow cytometric analysis of human immune cell composition in digested skin samples (left, per gram skin and 100,000 live cells), including expression of activation markers CD69 (middle column) and CD18 (right column). ΔMFI, median fluorescence intensity after subtraction of FMO control. (**E**) Quantification of murine immune cells in skin samples by flow cytometry (per gram skin and 100,000 live cells) (**F**) Quantification of human cytokines in plasma on day 13 using the LEGENDplex^TM^ Human Inflammation Panel 1. Symbols indicate individual mice; bars represent the mean. Statistical analyses were performed using Shapiro-Wilk normality test followed by unpaired t-test (gaussian distribution) or Mann-Whitney test (non-gaussian distribution). * p < 0.05; ** p < 0.01.

### PDK1 inhibition with BX-795 impairs myeloid cell activation and disease development

The PDK1/Akt signaling pathway, downstream of PI3K, plays a key role in modulating neutrophil function (*26, 27*). Moreover, in vitro kinome profiling of lesional skin from experimental PD mouse models identified PI3K as prominently activated (*20*) and a selective role of PI3K in FcγR-dependent neutrophil activation was demonstrated by a dose-dependent inhibition of ROS production upon treatment with a PI3K inhibitor (*25*). Importantly, several inhibitors of PI3K signaling were shown to successfully impair disease development in mouse models of PD (*18, 20, 58*). We therefore hypothesized that BX-795, a potent ATP-competitive inhibitor blocking PDK1 downstream of PI3K but upstream of Akt (*23, 59, 60*), would potently suppress the effector phase of EBA. In vitro, BX-795 was able to decrease ROS production in IgG-stimulated human primary cells, particularly neutrophils and to a minor extent also in classical monocytes (Fig. 4A). Secretion of inflammatory cytokines IL-1β, IL-8 and IL-18 was also reduced strongly supporting the potential of BX-795 to repress development of IgG-induced skin inflammation (Fig. 4B). Accordingly, in an ex vivo model, incubation of cryosections of human skin samples with human neutrophils in presence of EBA autoantibodies induced overall separation of the DEJ. This was prevented when cells were pre-treated with BX-795 (Fig. 4C). In line with these results, treatment of HIS mice with BX-795 (Fig. 5A) strongly reduced skin inflammation (Fig. 5B) and inflammatory infiltrations and epidermal thickness appeared to be decreased (Fig. 5C). Detailed analysis of human immune cells in the affected skin revealed reduced numbers of classical monocytes and neutrophils. Neutrophil activation was also impacted with lower expression of activation markers CD86 and CD18. Importantly, neutrophils were also found to produce less ROS (Fig. 5D). Unexpectedly, plasma concentrations of inflammatory cytokines were less affected but also subject to pronounced heterogeneity. Overall, these results not only establish the suitability of HIS mice in investigating EBA-like autoimmunity. They also, for the first time, show the therapeutic potential of small-molecule inhibition of PDK1 for its treatment.

**Fig. 4.**
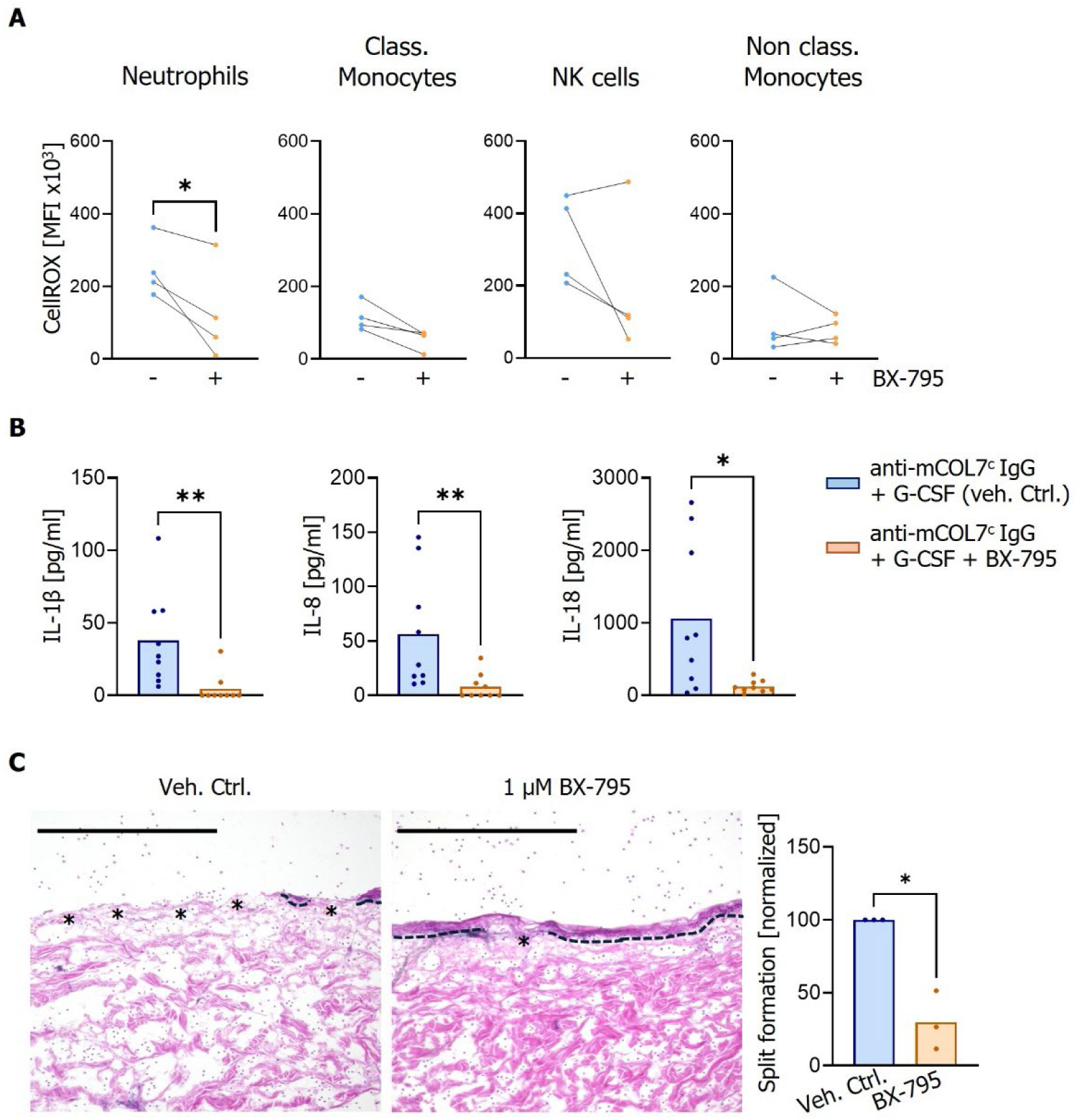
PDK1 kinase inhibition with BX-795 in human peripheral blood cells suppresses EBA effector functions in vitro. (**A**) Flow cytometric analysis of cytosolic ROS production in human peripheral blood cells after stimulation with COL7^c^-IgG immune complexes (IC) upon pre-treatment with 1µM BX-795 or vehicle control. n=4 independent experiments using cells from different donors were performed. (**B**) Quantification of secreted human cytokines in the supernatants of COL7^c^-IC stimulated human cells, comparing presence or absence of PDK1 kinase inhibitor BX-795, measured using the LEGENDplex^TM^ Human Inflammation Panel 1. (**C**) Induction of dermal-epidermal separation in cryosections of human skin was performed upon treatment of neutrophils with 1µM BX-795 compared to untreated control cells. Line symbols the dermal-epidermal junction, asterisks separation. Scale bar 500 µm. Statistical analyses employed the Shapiro-Wilk normality test. Depending on Gaussian distribution either paired t-test or Wilcoxon test was used. * p < 0.05; ** p < 0.01.

**Fig. 5.**
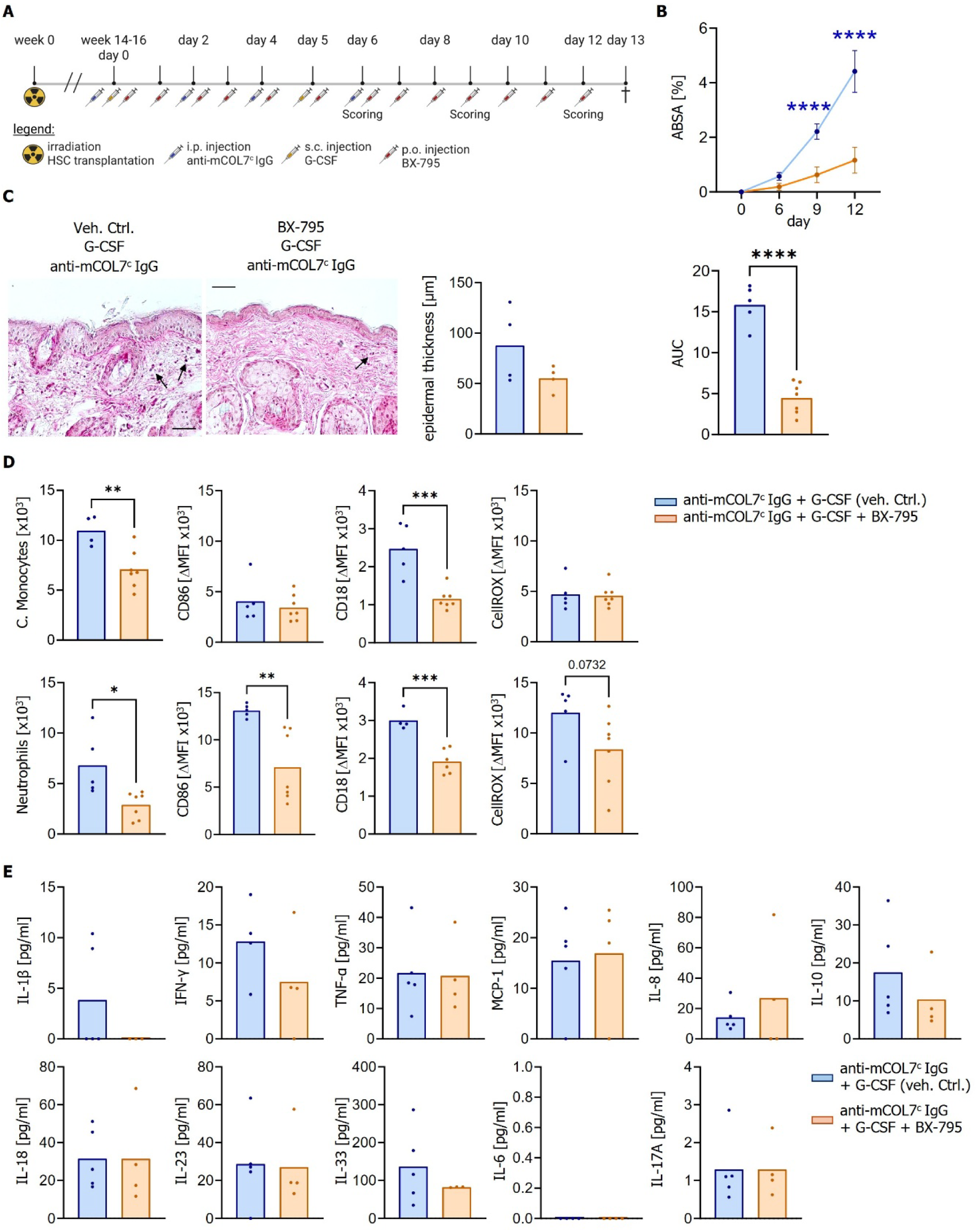
PDK1 kinase inhibition with BX-795 as therapeutic intervention in EBA-like autoimmunity in human immune system mice. (**A**) Schematic representation of the experimental approach for inducing an EBA-like phenotype in human immune system mice, combined with p.o. administration of 30 mg/kg BX-795 or vehicle as control. (**B**) Assessment of skin disease severity shown as % ABSA over time (left) and as AUC (right). (**C**) Histological analysis of skin sections using PAS staining for tissue structure and infiltrate evaluation (arrow) and quantification of epidermal thickness (in µm, right). Veh. Ctrl., vehicle control. (**D**) Human immune cell composition in digested skin samples (left, per gram skin and 100000 live cells), including expression of activation markers CD86 and CD18 (middle), and intracellular ROS production (right). C., classical. ΔMFI, median fluorescence intensity after subtraction of fluorescence minus one. (**E**) Quantification of human cytokines in plasma on day 6 using the LEGENDplex^TM^ Human Inflammation Panel 1. Symbols indicate individual mice and bars indicate means. Statistical analyses were performed using Shapiro-Wilk normality test. Depending on Gaussian distribution, samples were analyzed using unpaired t-test or Mann-Whitney test, respectively. *p < 0.05; **p < 0.01; ***p < 0.001; ****p <0.0001.

## DISCUSSION

The herein established experimental setup for EBA induction in human immune system mice allows significant and reproducible manifestation of disease depending on presence and activation of human immune effector cells and is therefore well-suited to assessing the clinical efficacy of therapeutic regimens targeting human immune cells. Importantly, human immune system mice not only feature major human leukocyte populations expressing human FcγRs which allows to investigate IgG activity in vivo. They also reflect the genetic complexity of the human immune system. Nevertheless, heterogeneity of humanization is a confounding factor considering the extent of disease development. We could partially address this by administration of recombinant human G-CSF which is known to support myeloid cell differentiation and maturation although this did not enhance disease to levels observed in patients. This was neither due to inefficient deposition of the autoantibodies at the DEJ (Suppl. Fig. S5) nor an impairment in cell infiltration into the skin but likely a consequence of still relatively limited numbers of crucial cell populations in HIS mice. Compared to humans, where neutrophils amount to 80 % of leukocytes in peripheral blood (*61*), this population only reached up to 5% of human cells (or on average 1-2 % of total mouse and human leukocytes) in HIS mice. Neutrophils were described as the most important myeloid effector cells in humans as well as mice (*5, 62*) and low numbers could explain reduced disease activity. Indeed, we observed a correlation of disease severity with the number of neutrophils, and to a lesser extent with classical monocytes (Fig. 2D). Thus, human immune system mice appear to have low, but sufficient numbers of functional myeloid cells able to drive disease manifestation.

In myeloid effector cells expressing human FcγRs (Suppl. Fig. S2) (*43*), the extent of cell activation and thus disease severity could be related to polymorphic variants of FcγRs previously described to affect IgG binding or signal transduction (*7, 35*). Therefore, we performed genotyping for common polymorphisms in FcγRIIa (histidine or arginine at amino acid position 131) (*63*), FcγRIIIa (valine or phenylalanine at amino acid position 158) (*64*), FcγRIIIb (NA1 or NA2 variant) (*65*) and FcγRIIb (isoleucine or threonine at amino acid position 232) (*66*). No enrichment of certain allelic variants could be observed (Suppl. Table 1) but the number of animals might currently be too small to draw definitive conclusions on this matter. In addition to FcγRs, the complement system is also involved in EBA development in mouse models (*4, 67*). The mouse strain used in this study carries a mutation in the gene encoding for complement factor C5. This mutation is characteristic for the NOD background and causes a premature STOP codon abrogating C5 expression (*68*). To the best of our knowledge, this has not been confirmed experimentally but one would consequently assume a deficiency in functional C5 in all strains with this background. Further experiments would be needed to confirm or exclude a contribution of C5 in NSG/FcRγ^-/-^ mice as an additional factor for disease activity in the present study.

The importance of human immune effector cells was confirmed by blocking of human FcγRs which resulted in strong reduction of disease severity. The combination of antibodies used in our study allows blocking of all human FcγRs but does not enable conclusions on the responsible effector cells. The current data suggest both neutrophils and classical monocytes to mediate skin inflammation based on their increased presence in skin infiltrates and activation. Identifying the responsible immune effector cell population will also be important for development of improved, more targeted therapies as opposed to current, broad immunosuppressing treatment schemes. Given the crucial role of FcγR-dependent immune cell activation for disease manifestation these could include targeting of molecules involved in intracellular signaling cascades. The upstream kinase Syk is a prominent example which was shown to be required for EBA manifestation in mice and patients (*62, 69, 70*) albeit the therapeutic activity of Syk inhibition in PD has not been explored to date. The inflammatory potential of the PI3K signaling pathway is equally well established (*71*). PI3K is also essential for FcγR-dependent neutrophil activation (*25*) and inhibition of PI3K was beneficial in a model of arthritis (*72*), a disease also driven by IgG autoantibodies. We therefore assessed the therapeutic capacity of BX-795, an inhibitor for PDK1 which is located downstream of PI3K (*23*), due to its direct association with disease-relevant neutrophil activation. BX-795 was originally developed for cancer therapy (*59*) but its anti-inflammatory capacity was meanwhile confirmed in vitro (*26, 73*) as well as in inflammatory mouse models in vivo (*60, 74*). Our results substantiate impaired neutrophil activation by BX-795 but also revealed reduced ROS production in monocytes and overall cytokine secretion. No adverse side effects were observed, and viability of myeloid cell populations appeared to be unaffected (Suppl. Fig. S8) despite systemic treatment of HIS mice over an extended period of time. This successful proof-of-concept for kinase inhibition in PD in HIS mice confirms the therapeutic potential and tolerability of kinase inhibition in autoimmunity and inflammation already precedented by successful application of JAK, Syk and BTK inhibitors in pemphigoid disease, inflammatory arthritis and immunothrombocytopenia, amongst others (*21, 75–77*).

Several limitations of our study should be acknowledged. First, disease was induced by passive transfer of a rabbit-derived COL7^c^-specific IgG as no recombinant antibody is currently available. Investigating disease pathomechanisms would greatly benefit from availability of human autoantibodies. Second, as described above, reconstitution of the human immune system is heterogenous affecting disease development, and the contribution of the complement system remains unclear. Third, this study does not address specificity of BX-795. While originally developed to target PDK1, BX-795 has meanwhile been shown to also impair TANK-binding kinase 1 (TBK1) and IĸB kinase (IKKɛ) which could contribute to its anti-inflammatory activity (*60*).

In the future, the present study could be followed up with inhibitors acting more downstream further increasing specificity related to distinct effector functions and cell types. Future studies should also address the suitability of applying this and related inhibitors topically to interfere with skin inflammation in a way that encourages patient compliance as well as reduces unwanted side effects. Together, these findings establish a uniquely human-relevant in vivo model and position PDK1 inhibition as a compelling new therapeutic avenue for pemphigoid diseases.

## MATERIAL AND METHODS

### Study Design

EBA is a rare disease, and access to patient material is limited. Therefore, the aim of this study was to establish an in vivo EBA model that enables investigation of disease pathogenesis and evaluation of novel therapeutic targets in the context of human cells and their FcγRs. We ultimately aimed to investigate the pre-specified hypothesis that PDK1 inhibition would impair EBA development in HIS mice. Rabbit IgG binding to human FcγRs and subsequent immune cell activation were assessed in vitro using human peripheral blood of five healthy volunteers. Control treatments were performed for each donor. For the in vivo study, HIS mice were generated by transplanting NSG/FcRγ^-/-^ mice with human HSCs. EBA was then induced in successfully humanized mice assigned to the treatment groups: (i) HIS mice injected with total rabbit IgG (negative control), (ii) HIS mice injected with anti-COL7^c^ specific rabbit IgG, (iii) HIS mice injected with anti-COL7^c^ specific rabbit IgG and recombinant G-CSF to increase myeloid cell numbers, (iv) HIS mice injected with anti-COL7^c^ specific rabbit IgG, recombinant G-CSF and human FcγR-blocking antibodies, and (v) HIS mice injected with anti-COL7^c^ specific rabbit IgG, recombinant G-CSF and daily treatment with BX-795 compared to vehicle control. Disease progression was monitored by clinical scoring of skin alterations over 13 days before endpoint-analysis. Overall, HIS mice from eight independent HSC donors were used. Each experimental group contained a minimum of five mice from at least two HSC donors to account for donor-specific variations. Animals were allocated to treatment groups via randomized selection while monitoring and accounting for heterogeneity in reconstitution (Suppl. Table 1). Investigators were not blinded to the treatment groups.

### In vitro assays

#### Binding of rabbit COL7^c^-specific IgG to human FcγRs

Chinese hamster ovary (CHO) cell lines stably expressing human FcγRI, -131Arg and -131His variants of FcγRIIa, FcγRIIb, -158F and -158V variants of FcγRIIIa, and -NA1 and -NA2 variants of FcγRIIIb (*41, 42, 78*) were used to assess binding of rabbit IgG. CHO cells expressing specific FcγRs or untransfected controls (200,000 cells per condition) were incubated with 10 µg/ml rabbit IgG or PBS for 1 h at 4°C under gentle shaking. Following incubation, bound IgG was detected by staining with a DyLight594-conjugated anti-rabbit IgG (0.5µg/mL; BioLegend) followed by acquisition of median fluorescence intensity (MFI) on a CytoFLEX S flow cytometer (Beckman Coulter).

In addition, binding to human primary peripheral blood leukocytes (PBL) was investigated using whole blood from five healthy volunteers. Erythrocytes were lysed by deionized water for 20 s at room temperature and binding assays were subsequently performed as described above. Leukocyte subpopulations were identified by staining with surface marker-specific fluorescence-conjugated antibodies: PE/Cy7 anti-CD19 (SJ25C1), PerCP anti-CD3 (UCHT1), PE anti-CD33 (WM53), Brilliant Violet 510 anti-CD14 (M5E2), FITC anti-CD56 (MEM188) and APC/Fire 750 anti-CD45 (2D1) (all BioLegend). MFI of bound rabbit IgG were detected using a DyLight594-conjugated anti-rabbit IgG (Poly4064, BioLegend) at 0.5 μg/mL and samples were acquired using a FACSCanto II flow cytometer (BD).

Cell populations were gated as follows: Aggregates were excluded by forward scatter area/height ratio (FSC-A/FSC-H); dead cells excluded by DAPI staining; leukocytes identified by CD45 expression. Neutrophils were gated as side light scatter (SSC) high and CD14^-^; classical monocytes as SSC-intermediate, CD14^+^. Within the CD14^−^SSC^low^ population, B (CD19^+^) and T cells (CD3^+^) were identified. NK cells were distinguished by CD56 expression, and non-classical monocytes were gated as CD33^+^ among remaining cells.

Analysis of flow cytometry data was performed with FlowJo V10. Background fluorescence intensity as observed in fluorescence minus one (FMO) control was subtracted.

### Leukocyte activation by rabbit COL7^c^-specific IgG

Production of reactive oxygen species (ROS) is a key effector function in the pathogenesis of EBA (*5*). Human leukocytes, purified as described above, were stimulated with immobilized COL7^c^/anti-COL7^c^ IgG immune complexes (IC) at concentrations of 10 µg/ml (COL7^c^) and 2 µg/ml (anti- COL7^c^ IgG) in a 96-well flat-bottom plate for 30 min at 37 °C and 5 % CO_2_. To verify the effect of PDK1 inhibition, human cells were treated with BX-795 (1 µM in 0.1 % DMSO) or buffer (vehicle control) for 5 min at room temperature prior to stimulation. Stimulated cells were then transferred to a 96-well V-bottom plate and stained with 5 µM CellROX Deep Red reagent (Thermo Fisher Scientific) for 30 min at 37 °C. During the last 15 min of incubation, fluorescently labelled antibodies against cell-type-specific surface markers were added to identify leukocyte subpopulations. The following antibodies were used: PerCp anti-CD3 (UCHT1), BV510 anti-CD14 (M5E2), Spark Blue 550 anti-human CD45 (2D1), BV650 anti-CD19 (HIB19), BV605 anti-CD16 (3G8), FITC anti-CD56 (MEM188), PE anti-CD33 (WM53) and PE/Cy7 anti-CD66b (g10F5) (all BioLegend). Cell populations were gated as follows: Aggregates excluded by forward scatter area/height ratio (FSC-A/FSC-H); dead cells excluded by Zombie NIR staining; human leukocytes were identified with human CD45. Within the human cells, B cells were identified as CD19+ and T cells as CD3+; NK cells were gated as CD56+ in the CD19-CD3- population; within the CD56- gate, neutrophils were identified as CD66b+ and remaining myeloid cells were CD33+. Classical monocytes (CD14+CD16-) and non-classical monocytes (CD14-CD16+) were gated within CD33+ cells. Data acquisition was performed on a Northern Lights flow cytometer (Cytek). Intracellular ROS production was quantified as the median fluorescence intensity (MFI) of the APC channel representing the CellROX signal for each cell population. Seven independent human donors were analyzed with unstimulated cells serving as negative controls.

Cell culture supernatants from unstimulated and stimulated (treated with BX-795 or vehicle control) cells were collected and stored at -80°C until cytokine profiling was performed using the LEGENDPlex^TM^ Human Inflammation Panel 1 (BioLegend) according to the manufactureŕs instructions. Data was acquired on the Northern Lights flow cytometer (Cytek) and analyzed using Qognit software to determine cytokine concentrations relative to standard curves.

### Cryosection assay of human skin

To assess the capacity of BX-795 to block split formation ex vivo, cryosection assays were performed as described before with modifications (*5*). Blood was obtained from healthy human volunteers. Neutrophils were isolated by density gradient centrifugation and erythrocytes were lysed with 20 ml cold 0.2 % (w/v) NaCl solution for 30 s at RT and stopped with an equal volume of cold 1.6 % (w/v) NaCl solution. After centrifugation, the pellet was resuspended in 10 ml RPMI 1640 with L-glutamine and stored on ice and the cell number adjusted to 5x10^7^. Meanwhile, 6 µm thick cryosections of human breast skin biopsies were thawed and 30 µl recombinant human anti-COL7 IgA2 were applied per section. The slides were incubated in a humid chamber for 1 h at 37 °C followed by washing in PBS for 5 min at RT. The slides were each covered with another slide using adhesive tape leaving space in between for approximately 500 µL per section. BX-795 was added to the cell suspension at a final concentration of 1 µM. 500 µL cell-inhibitor suspension were added per section and the slides were incubated in a humid chamber for 3 h at 37 °C. Subsequently, the additional slides were removed, and the slides were washed in PBS until the dermal-epidermal junction was visible. The sections were stained for H&E using standard protocols. Images were taken at a BZ-9000E series microscope (Keyence). Split formation was evaluated in relation to the total length of the dermal-epidermal junction using ImageJ (*79*).

### In vivo assays Mice

NSG-FcRγ^−/−^ mice were generated by backcrossing SCID, γc-deficient, and FcRγ^−/−^ mice onto the NOD background for at least 10 generations followed by intercrossing (*80, 81*). Due to their immunocompromised status, NSG-FcRγ^−/−^ mice were provided with acidified drinking water (pH 3.0) to reduce the risk of bacterial infection. All experiments were conducted using male and female mice aged between 14 to 16 weeks. Mice were housed under specific-pathogen-free conditions in isolated ventilated cages at the animal facility of Friedrich-Alexander-University Erlangen-Nuremberg, in compliance with the rules and regulations of the German animal welfare. All animal experiments were approved by the government of lower Franconia.

### Generation of human immune system HIS mice

HIS mice were generated as described (*43, 44, 80–83*). In brief, newborn NSG-FcRγ^−/−^ mice were irradiated with a dose of 1.4 Gy and intravenously injected with 20,000–40,000 human HSCs 6-8 h post-irradiation. HSCs were isolated from umbilical cord blood with written informed consent from donors as previously described. The efficiency of human immune system reconstitution was assessed 12 to 14 weeks after transplantation by flow cytometry analysis of human cell subsets in whole blood. The following antibodies were used: cFluor V420 anti-CD3 (SK7), cFluor V450 anti-CD14 (M5E2), cFluor V547 anti-human CD45 (HI30), cFluor BYG710 anti-CD19 (HIB19), cFluor R668 anti-CD16 (3G8) and cFluor R720 anti-CD56 (5.1H11), all from Cytek® cFluor® TBMNK Kit. Additionally, BV650 anti-murine CD45 (A20), BV605 anti-CD33 (P67.6) and PE/Cy7 anti-CD66b (G10F5) (BioLegend) were used. Cell populations were gated as follows: Aggregates were excluded by FSC-A/FSC-H ratio; dead cells excluded by Zombie NIR staining; human and murine leukocytes were separated with human CD45 against murine CD45. Within the human cells, B cells were identified as CD19^+^ and T cells as CD3^+^; NK cells were gated as CD56^+^ in the CD19^-^CD3^-^population; within the CD56^-^ gate, neutrophils were identified as CD66b^+^ and remaining myeloid cells were CD33^+^. Classical monocytes (CD14^+^CD16^-^) and non-classical monocytes (CD14^-^CD16^+^) were gated within the CD33^+^ population. Data was aquired on a Cytek Northern Lights flow cytometer and analyzed with FlowJo V10.

### Isolation of genomic DNA and FcγR genotyping

Genotyping of cord blood samples for FcγRIIa (131-Arg/His), FcγRIIIa (158-Phe/Val) and FcγRIIb (232-Ile/Thr transmembrane and 386G/C promoter polymorphism) alleles was done as described before (*43, 82*). Briefly, 250 µL of umbilical cord blood was collected and stored at -20°C. Genomic DNA was isolated with the „QiaAmp DSP Blood MiniKit “(Qiagen) following the instructions of distributor. FcγRIIa and FcγRIIIa genotyping were carried out with a two-step PCR protocol with an allele-specific nested PCR approach (*82*) and FcγRIIb genotyping with a two-step PCR protocol amplifying the promoter or transmembrane region, respectively, followed by sequencing (*83*). For FcγRIIIb genotyping, a 2.7kb product was amplified using (forward primer: 5′-cagaagatctcccaaaggctg-3′; reverse primer: 5′-gacccgaaacgtaataagagc-3′). Followed by purification via gel electrophoresis. The presence of five single nucleotide polymorphisms (*84*) was checked upon sequencing.

### EBA induction in HIS mice

A passive antibody transfer model was used to induce an EBA-like phenotype in HIS mice. To obtain anti-COL7^c^ rabbit IgG, New Zealand white rabbits were s.c. immunized with 250 mg murine von Willebrand factor A-like domain 2 protein suspended in CFA. The animals were boosted three times (at 13-day intervals) with the same protein preparation in IFA, and immune sera were collected (Eurogentec, Belgium). IgG from rabbit sera was purified using protein G affinity chromatography followed by affinity purification of anti-COL7^c^ IgG (*31, 32*). 100 µg of rabbit COL7^c^-specific IgG per 25 g body weight was injected i.p. on days 0, 2, 4 and 6. Total rabbit IgG (depleted for anti-COL7^c^-specific IgG) served as control. To increase human myeloid cell numbers in the blood, 50µg of recombinant human G-CSF was administered s.c. on days 0 and 5 following anti-COL7^c^-specific IgG injections. To assess the role of human FcγRs, selected animals received additional treatments with 100 µg of monoclonal antibodies targeting human FcγRI (10.1), FcγRIIa/b (IV.3) and FcγRIIIa/b (3G8) on days 0, 3, 6, 9 and 12. For evaluating the kinase PDK1 as a therapeutic target, HIS mice were treated orally with the small-molecule inhibitor BX-795 (Selleckchem) at 30 mg/kg daily. The inhibitor was dissolved in 5% DMSO, 30% PEG-300, 0,5% Tween-80 and 64,5% water with the buffer solution also used as vehicle control. Disease progression was monitored by clinical scoring of skin lesions on days 6, 9 and 12 post initial antibody administration. On day 13, mice were sacrificed, and skin samples were collected from the trunk after removing the fur.

Blood samples were obtained on days 0, 3, 6, 9 and 12, plasma isolated by centrifugation and stored at -80 °C until cytokine profiling was performed using the LEGENDPlex^TM^ Human Inflammation Panel 1 and Mouse Inflammation Panel for quantification of human and murine cytokines, respectively. Data was acquired on the Northern Lights flow cytometer (Cytek) and analysed using Qognit software to determine cytokine concentrations relative to standard curves.

### Generation of single-cell suspensions of skin samples

Skin samples from HIS mice were enzymatically digested to obtain single-cell suspensions. Samples were finely minced and incubated with the enzymatic mix from the Human Whole Skin Dissociation kit (Miltenyi Biotec). Up to 0.06 g of skin tissue was digested per recommended enzyme volume. Digestion was performed for 2 hr at 37 °C and 500 rpm. The enzymatic reaction was stopped by adding ice-cold DMEM with stable glutamine, 10% fetal bovine serum (FBS), 10 mM HEPES, 1% Penicillin/Streptavidin, 1mM sodium pyruvate, and 1× non-essential amino acids. The resulting cell suspensions were sequentially filtered through 70 µm and 40 µm cell strainers, centrifuged at 300x g for 10 min at 10 °C and subsequently processed for flow cytometry analysis. Human immune cell subsets were identified as described above with addition of PE-Dazzle594 anti-CD18 (TS1/18), PE anti-CD69 (FN50) and Pe-Cy5 anti-CD86 (IT2.2) to analyze the expression of human activation markers on the cell surfaces. For identification of murine immune cells, the following antibodies were used: BV650 anti-murine CD45 (A20), BV421 anti-CD11b (M1/70), Spark Blue 550 anti-Ly6G (1A8), PE-Dazzle594 anti-Ly6C (HK1.4). Cell populations were gated as follows: Aggregates excluded by forward scatter area/height ratio (FSC-A/FSC-H); dead cells excluded by Zombie NIR staining; human and murine leukocytes were separated with human CD45 against murine CD45. Murine myeloid cells were identified as CD11b^+^. Murine neutrophils were gated as Ly6G^+^ and classical monocytes as Ly6C^+^. Ly6G^-^Ly6C^-^ cells were identified as non-classical monocytes. Data was aquired on a Northern Lights flow cytometer (Cytek) and analyzed with FlowJo V10.

### Histological analysis of skin samples

Skin samples were embedded in paraffin for periodic acid-Schiff (PAS) staining and in O.C.T compound medium for immunofluorescence microscopy. Samples were fixed overnight at 4 °C in 4 % paraformaldehyde before paraffin embedding using a tissue processor (STP 120, Thermo Scientific). Paraffin-embedded samples were then sectioned into 5 µm slices using a MEDITE A550 microtome, followed by overnight fixation at 37 °C prior to PAS staining. The staining was performed as follows: All sections were first deparaffinized by submersion in xylene for 20 min, followed by successive treatments of 100 % ethanol, 96 % ethanol and 75 % ethanol, each for 2 min. The tissue was rehydrated in deionized water for 5 min. Next, they were treated with a freshly made 0.5 % periodic acid solution for 5 min and washed two times with deionized water. The sections were subsequently treated with Schiff’s reagent for 15 min and then washed under running tap water for 5 min. Finally, the nuclei were stained by submersing the sections in haematoxylin for 4 min and washed under running tap water for 5 min once more. After staining, the slides were treated with ascending ethanol concentrations consisting of 75 % ethanol, 96 % ethanol and 100 % ethanol for each 1 min. The slides were submersed in xylene for at least 5 min, after which they were mounted with Entellan™ mounting medium. Stained sections were analyzed using a Axiovert 200 microscope (Zeiss). Epidermal thickness at three locations each was quantified for five exemplary images per mouse and mean thickness calculated.

For immunofluorescence microscopy, samples embedded in O.C.T medium were flash-frozen on dry ice and stored at -80 °C. 5 µm cryosections were cut using a cryotome, fixed for 10 min in ice cold acetone, and washed three times in 1x PBS. Immunofluorescence staining was performed to visualize rabbit IgG deposits along the DEJ. Nonspecific binding sites were blocked with 5 % goat serum before incubation with DyLight594-conjugated anti-rabbit IgG antibody (BioLegend) for 1 hr at room temperature in a humidified chamber. Stained sections were analyzed using a Zeiss Axiovert 200 microscope.

## Ethics statement

Purification of HSCs from umbilical cord blood as well as application of primary leukocytes purified from peripheral blood were approved by the Ethics Committee of FAU Erlangen and Lübeck. The studies were conducted in accordance with the local legislation and institutional requirements. The participants provided their informed consent.

## Statistics

Statistical analysis was performed using the software GraphPad Prism, version 9. Data sets were tested for normal distribution using the Shapiro-Wilk normality test. Depending on sample distribution, paired t-tests, Wilcoxon tests, Ordinary one-way ANOVA, Kruskal-Wallis tests, unpaired t-tests and Mann-Whitney tests were used. Tests and significance p values are described in the respective figure legend.

## List of Supplementary materials

Table S1. Overview of HIS mice used in study.

Fig. S1. Humanization efficiency of NSG/FcRγ^-/-^ mice.

Fig. S2. Gating strategies for identification of human leukocyte populations in HIS mouse blood and skin.

Fig. S3. EBA-like autoimmunity in irradiated non humanized NSG/FcRγ^-/-^ mice.

Fig. S4. Human G-CSF is increasing and maturing human myeloid cells in the blood of HIS mice.

Fig. S5. Deposition of rabbit anti-COL7^c^ IgG in skin of HIS mice.

Fig. S6. Human cytokines in the plasma of HIS mice after EBA induction.

Fig. S7. Murine cytokines in the plasma of HIS mice after EBA induction.

Fig. S8. Viability of human cells treated with BX-795 in vitro and in vivo.

## Acknowledgements

We thank Elza Gareus, Marta Delogu, Fabien Isbrecht, Georg Kaiser and Azaliya Yabbarova for expert technical assistance and Sripriya Murthy for providing rabbit anti-COL7 IgG and control IgG.

## Author contributions

Conceptualization: KB, AL

Investigation: LV, KM, SW, AS, MS, LA, EU, LH, MP Funding acquisition: KB, AL

Supervision: FN, KB, AL

Writing – original draft: LV, KB, AL

Writing – review & editing: LV, KM, SW, AS, MS, LA, EU, LH, MP, FN, KB, AL

## Data and materials availability

All data associated with this study are present in the paper or the Supplementary Materials. Further data that support the findings of this study are available from the corresponding author upon reasonable request.

