## Supplementary Material for "A human immune system mouse model for preclinical evaluation of therapies in pemphigoid disease"

### Suppl. Table 1

| Fig. | treatment | HIS mouse | | | FcgR polymorphism | | | | | hCD45+ cells [% of live] | % of hCD45+ | | | | | | |
|---|---|---|---|---|---|---|---|---|---|---|---|---|---|---|---|---|---|
| | | mouse # | HSC donor # | sex | FcγRIIa 131H/R | FcγRIIIa 158F/V | FcγRIIb 232I/T | FcγRIIb 386G/C | FcgRIIIb NA1/NA2 | | B cells | T cells | NK cells | class. Monocytes | int. Monocytes | non class. Monocytes | neutrophils |
| 2 | rabbit IgG | **1** | A | m | H/R | F/V | I/I | PP-wt | NA2 | **9,8** | 73,6 | 0,2 | 1,0 | 9,6 | 1,7 | 0,5 | 2,4 |
| 2 | rabbit IgG | **2** | D | m | H/H | F/V | I/I | PP-wt | NA1/2 | **9,7** | 84,7 | 4,9 | 0,1 | 3,4 | 0,0 | 0,0 | 0,1 |
| 2 | rabbit IgG | **3** | D | f | H/H | F/V | I/I | PP-wt | NA1/2 | **40,1** | 88,6 | 8,3 | 0,1 | 1,1 | 0,0 | 0,1 | 0,1 |
| 2 | rabbit IgG | **4** | D | m | H/H | F/V | I/I | PP-wt | NA1/2 | **31,9** | 95,9 | 1,1 | 0,3 | 1,0 | 0,0 | 0,4 | 0,1 |
| 2 | rabbit IgG | **5** | E | m | H/R | F/V | I/I | PP-wt | NA1 | **8,4** | 6,3 | 72,4 | 1,6 | 5,8 | 0,0 | 1,5 | 5,1 |
| 2 | rabbit IgG | **6** | E | m | H/R | F/V | I/I | PP-wt | NA1 | **41,7** | 2,6 | 78,7 | 0,3 | 6,6 | 0,0 | 2,1 | 5,1 |
| 2 | anti-mCOL7c IgG | **7** | A | m | H/R | F/V | I/I | PP-wt | NA2 | **7,0** | 40,0 | 26,4 | 4,4 | 5,1 | 1,5 | 7,9 | 11,9 |
| 2 | anti-mCOL7c IgG | **8** | A | f | H/R | F/V | I/I | PP-wt | NA2 | **9,8** | 66,5 | 17,4 | 1,0 | 2,0 | 0,6 | 2,9 | 1,6 |
| 2 | anti-mCOL7c IgG | **9** | A | m | H/R | F/V | I/I | PP-wt | NA2 | **26,9** | 44,5 | 31,8 | 4,6 | 3,2 | 2,2 | 8,1 | 1,6 |
| 2 | anti-mCOL7c IgG | **10** | B | m | H/R | F/V | I/I | PP-wt | n.d. | **27,1** | 88,1 | 2,8 | 0,5 | 4,1 | 1,1 | 0,3 | 0,1 |
| 2 | anti-mCOL7c IgG | **11** | B | m | H/R | F/V | I/I | PP-wt | n.d. | **60,9** | 67,1 | 23,6 | 1,7 | 3,0 | 1,4 | 1,1 | 0,5 |
| 2 | anti-mCOL7c IgG | **12** | B | f | H/R | F/V | I/I | PP-wt | n.d. | **17,0** | 88,8 | 0,6 | 0,8 | 5,1 | 0,3 | 0,2 | 0,1 |
| 2 | anti-mCOL7c IgG | **13** | C | m | H/R | F/F | T/T | PP-wt | NA1/2 | **5,6** | 50,8 | 33,4 | 0,3 | 3,7 | 0,0 | 0,9 | 0,5 |
| 2 | G-CSF + anti-mCOL7c IgG | **14** | A | m | H/R | F/V | I/I | PP-wt | NA2 | **19,3** | 32,7 | 23,8 | 1,9 | 4,5 | 0,7 | 3,0 | 12,3 |
| 2 | G-CSF + anti-mCOL7c IgG | **15** | B | f | H/R | F/V | I/I | PP-wt | n.d. | **40,7** | 66,0 | 22,1 | 1,7 | 2,8 | 2,4 | 0,6 | 0,3 |
| 2 | G-CSF + anti-mCOL7c IgG | **16** | B | f | H/R | F/V | I/I | PP-wt | n.d. | **21,7** | 61,4 | 9,1 | 0,3 | 4,2 | 1,4 | 1,2 | 0,6 |
| 2 | G-CSF + anti-mCOL7c IgG | **17** | C | f | H/R | F/F | T/T | PP-wt | NA1/2 | **8,0** | 53,4 | 35,0 | 1,5 | 2,6 | 0,0 | 0,6 | 0,3 |
| 2 | G-CSF + anti-mCOL7c IgG | **18** | C | m | H/R | F/F | T/T | PP-wt | NA1/2 | **7,3** | 14,7 | 69,9 | 0,2 | 5,8 | 0,0 | 0,2 | 0,6 |
| 2 | G-CSF + anti-mCOL7c IgG | **19** | D | m | H/H | F/V | I/I | PP-wt | NA1/2 | **15,5** | 89,3 | 4,5 | 0,2 | 2,5 | 0,0 | 0,2 | 0,1 |
| 3 | G-CSF + anti-mCOL7c IgG | **20** | C | f | H/R | F/F | T/T | PP-wt | NA1/2 | **5,3** | 4,0 | 87,0 | 0,0 | 2,8 | 0,0 | 0,2 | 0,1 |
| 3 | G-CSF + anti-mCOL7c IgG | **21** | C | f | H/R | F/F | T/T | PP-wt | NA1/2 | **5,6** | 10,5 | 81,2 | 0,3 | 2,4 | 0,0 | 0,1 | 0,5 |
| 3 | G-CSF + anti-mCOL7c IgG | **22** | D | f | H/H | F/V | I/I | PP-wt | NA1/2 | **35,7** | 95,6 | 0,1 | 0,3 | 1,3 | 0,0 | 0,3 | 0,2 |
| 3 | G-CSF + anti-mCOL7c IgG | **23** | D | m | H/H | F/V | I/I | PP-wt | NA1/2 | **38,4** | 89,7 | 5,0 | 0,5 | 1,5 | 0,0 | 0,3 | 0,2 |
| 3 | G-CSF + anti-mCOL7c IgG | **24** | D | m | H/H | F/V | I/I | PP-wt | NA1/2 | **36,9** | 90,3 | 6,8 | 0,1 | 1,0 | 0,0 | 0,1 | 0,1 |
| 3 | anti-hFcγR + G-CSF + anti-mCOL7c IgG | **25** | A | m | H/R | F/V | I/I | PP-wt | NA2 | **23,1** | 41,1 | 14,7 | 3,8 | 5,8 | 2,2 | 7,8 | 4,7 |
| 3 | anti-hFcγR + G-CSF + anti-mCOL7c IgG | **26** | A | f | H/R | F/V | I/I | PP-wt | NA2 | **51,2** | 54,8 | 21,6 | 2,1 | 2,4 | 1,9 | 5,4 | 5,0 |
| 3 | anti-hFcγR + G-CSF + anti-mCOL7c IgG | **27** | B | m | H/R | F/V | I/I | PP-wt | n.d. | **31,5** | 51,2 | 41,4 | 0,6 | 3,6 | 0,9 | 0,2 | 0,2 |
| 3 | anti-hFcγR + G-CSF + anti-mCOL7c IgG | **28** | B | m | H/R | F/V | I/I | PP-wt | n.d. | **24,1** | 62,4 | 32,2 | 0,6 | 1,2 | 0,3 | 0,4 | 0,1 |
| 3 | anti-hFcγR + G-CSF + anti-mCOL7c IgG | **29** | D | f | H/H | F/V | I/I | PP-wt | NA1/2 | **22,5** | 95,6 | 0,0 | 0,2 | 1,4 | 0,0 | 0,2 | 0,1 |
| 3 | anti-hFcγR + G-CSF + anti-mCOL7c IgG | **30** | D | m | H/H | F/V | I/I | PP-wt | NA1/2 | **39,8** | 90,8 | 5,8 | 0,3 | 1,0 | 0,0 | 0,2 | 0,1 |
| 3 | anti-hFcγR + G-CSF + anti-mCOL7c IgG | **31** | D | f | H/H | F/V | I/I | PP-wt | NA1/2 | **29,6** | 89,0 | 7,5 | 0,1 | 1,0 | 0,0 | 0,1 | 0,1 |
| 5 | vehicle control | **32** | F | m | H/H | F/V | I/T | PP-wt | NA1/2 | **11,8** | 91,5 | 0,0 | 0,2 | 5,5 | 0,2 | 0,0 | 1,1 |
| 5 | vehicle control | **33** | F | m | H/H | F/V | I/T | PP-wt | NA1/2 | **19,1** | 89,7 | 0,0 | 0,1 | 7,7 | 0,2 | 0,0 | 1,2 |
| 5 | vehicle control | **34** | F | f | H/H | F/V | I/T | PP-wt | NA1/2 | **22,8** | 88,7 | 0,0 | 0,2 | 8,6 | 0,1 | 0,0 | 1,1 |
| 5 | vehicle control | **35** | H | m | R/R | F/F | I/I | PP-wt | NA2 | **16,4** | 93,3 | 0,4 | 0,1 | 3,9 | 0,1 | 0,1 | 0,4 |
| 5 | vehicle control | **36** | H | m | R/R | F/F | I/I | PP-wt | NA2 | **38,3** | 41,4 | 20,5 | 1,2 | 11,8 | 0,1 | 0,1 | 12,1 |
| 5 | BX-795 | **37** | F | f | H/H | F/V | I/T | PP-wt | NA1/2 | **13,8** | 85,3 | 0,0 | 0,2 | 9,5 | 0,1 | 0,0 | 4,9 |
| 5 | BX-795 | **38** | F | m | H/H | F/V | I/T | PP-wt | NA1/2 | **40,6** | 89,5 | 0,0 | 0,2 | 7,3 | 0,1 | 0,2 | 2,2 |
| 5 | BX-795 | **39** | F | f | H/H | F/V | I/T | PP-wt | NA1/2 | **42,9** | 87,7 | 0,1 | 0,2 | 8,7 | 0,0 | 0,1 | 2,8 |
| 5 | BX-795 | **40** | G | m | H/R | F/V | I/I | PP-wt | NA2 | **36,0** | 81,7 | 0,0 | 0,8 | 14,7 | 0,3 | 0,0 | 1,7 |
| 5 | BX-795 | **41** | H | f | R/R | F/F | I/I | PP-wt | NA2 | **32,6** | 79,6 | 6,7 | 4,3 | 2,5 | 0,1 | 0,2 | 1,9 |
| 5 | BX-795 | **42** | H | m | R/R | F/F | I/I | PP-wt | NA2 | **13,0** | 37,5 | 12,0 | 3,7 | 1,9 | 0,2 | 0,2 | 34,2 |
| 5 | BX-795 | **43** | H | m | R/R | F/F | I/I | PP-wt | NA2 | **33,2** | 66,3 | 18,7 | 1,8 | 4,6 | 0,0 | 0,0 | 5,0 |

Suppl. Table 1. **Overview of HIS mice used in study**

The study consisted of a total 43 mice generated from 8 HSC donors. Male and female animals were used. Humanization (hCD45$^{+}$) and composition of the human leukocytes were determined in peripheral blood before initiation of EBA. Mice were randomized on treatment groups ensuring comparable immune cell reconstitution between groups. Polymorphic variations in FcγRs were determined by allele-specific nested PCR (FcγRIIa-131H/R, FcγRIIIa-158F/V) or sequencing (FcγRIIb-232I/T, FcγRIIb-386G/C, FcγRIIIb-NA1/2). Experimental animals are sorted based on the figure they are shown in and treatment groups. Despite pronounced heterogeneity in human immune system reconstitution, no significant differences were present between treatment groups which could have impacted on differences observed in disease development. n.d. not determined

### Suppl. Figure 1

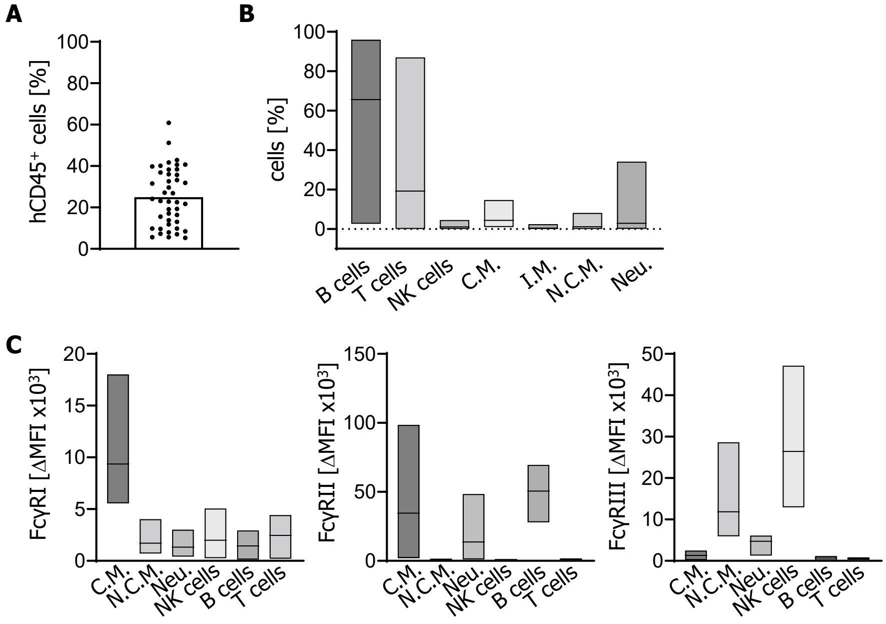


Fig. S1. **Humanization efficiency of NSG/FcRγ$^{-/-}$ mice.**
(**A**) Human CD45$^+$ cells in the blood of humanized NSG/FcRγ$^{-/-}$ mice. Cells are shown as % of live cells. **(B**) Development of all major human immune cell populations in the blood is shown as % of human CD45$^+$ cells. (**C**) Expression of human FcγRs on human immune cells as ΔMFI x10$^3$. C.M., classical monocytes, I.M., intermediate mmonocytes, N.C.M., non classical monocytes, Neu., neutrophils.

### Suppl. Figure 2

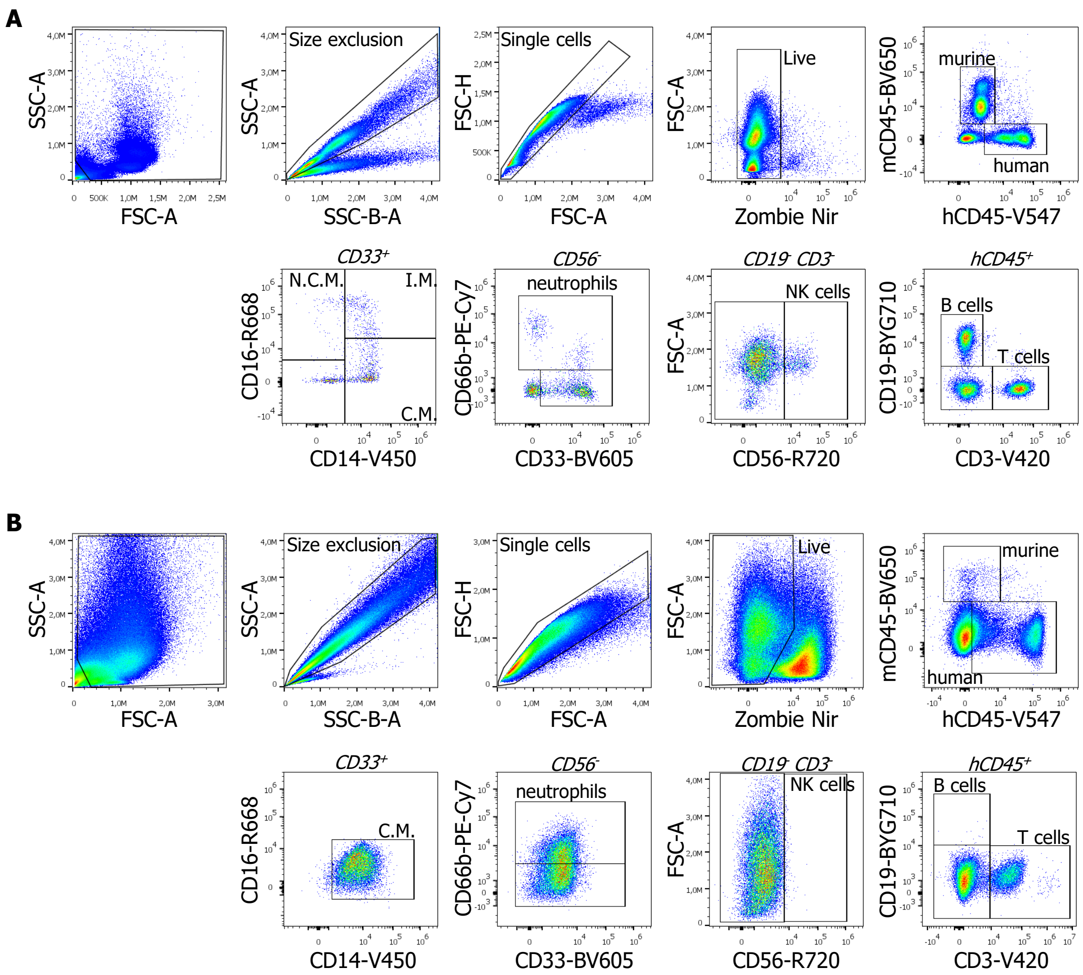


Fig. S2 **Gating strategies for identification of human leukocyte populations in HIS mouse blood and skin.**
(**A**) Gating strategy for identification of human leukocyte populations in HIS mouse blood. (**B**) Gating strategy for identification of human leukocyte populations in HIS mouse skin. Cell populations were gated as follows: aggregates were excluded by FSC-A/FSC-H ratio; dead cells excluded by Zombie NIR staining; human and murine leukocytes were separated with human CD45 against murine CD45. Within the human cells, B cells were identified as $CD19^+$ and T cells as $CD3^+$; NK cells were gated as $CD56^+$ in the $CD19^-CD3^-$ population; within the $CD56^-$ gate, neutrophils were identified as $CD66b^+$ and remaining myeloid cells were $CD33^+$. Classical monocytes ($CD14^+CD16^-$) and non-classical monocytes ($CD14^-CD16^+$) were gated within the $CD33^+$ gate.

### Suppl. Figure 3

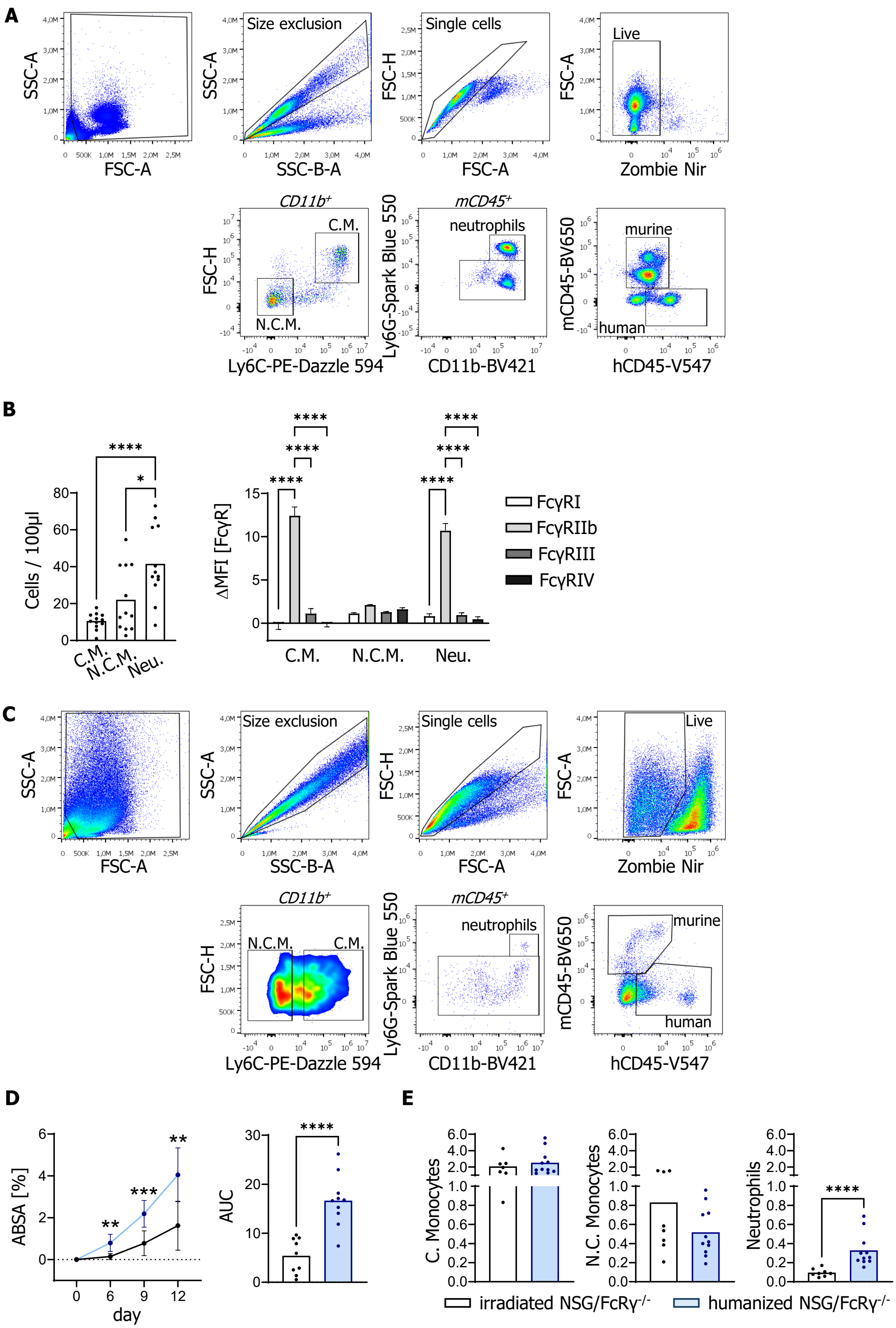


Fig. S3. **EBA-like autoimmunity in irradiated non humanized NSG/FcRγ$^{-/-}$ mice.**
(**A**) Gating strategy of murine leukocyte populations in blood of NSG/FcRγ$^{-/-}$ mice. (**B**) Murine immune cell composition (left) and FcγRs expression (right) in the blood of irradiated, non humanized NSG/FcRγ$^{-/-}$ mice (cells x10$^3$ per 100 µl blood). C.M., classical monocytes, N.C.M., non classical monocytes, Neu., neutrophils. **(C)** Gating strategy of murine leukocyte populations in skin of NSG/FcRγ$^{-/-}$ mice. **(D)** Assessment of skin disease severity shown as affected body surface area (ABSA, %) over time (left) and as area under the curve (AUC, right) of non humanized mice compared to G-CSF treated humanized mice. **(E)** Quantification of murine immune cells in skin samples by flow cytometry (per 1 g skin and 100,000 live cells). C., Classical, N.C., non classical. In (B) samples were analysed using Tukey's multiple comparisons test. In (D,E) Statistical analyses were performed using Shapiro-Wilk normality test. Depending on Gaussian distribution, samples were analysed using unpaired t-test or Mann-Whitney test, respectively. *$p < 0.05$; **$p < 0.01$; ***$p < 0.001$; ****$p < 0.0001$.

### Suppl. Figure 4

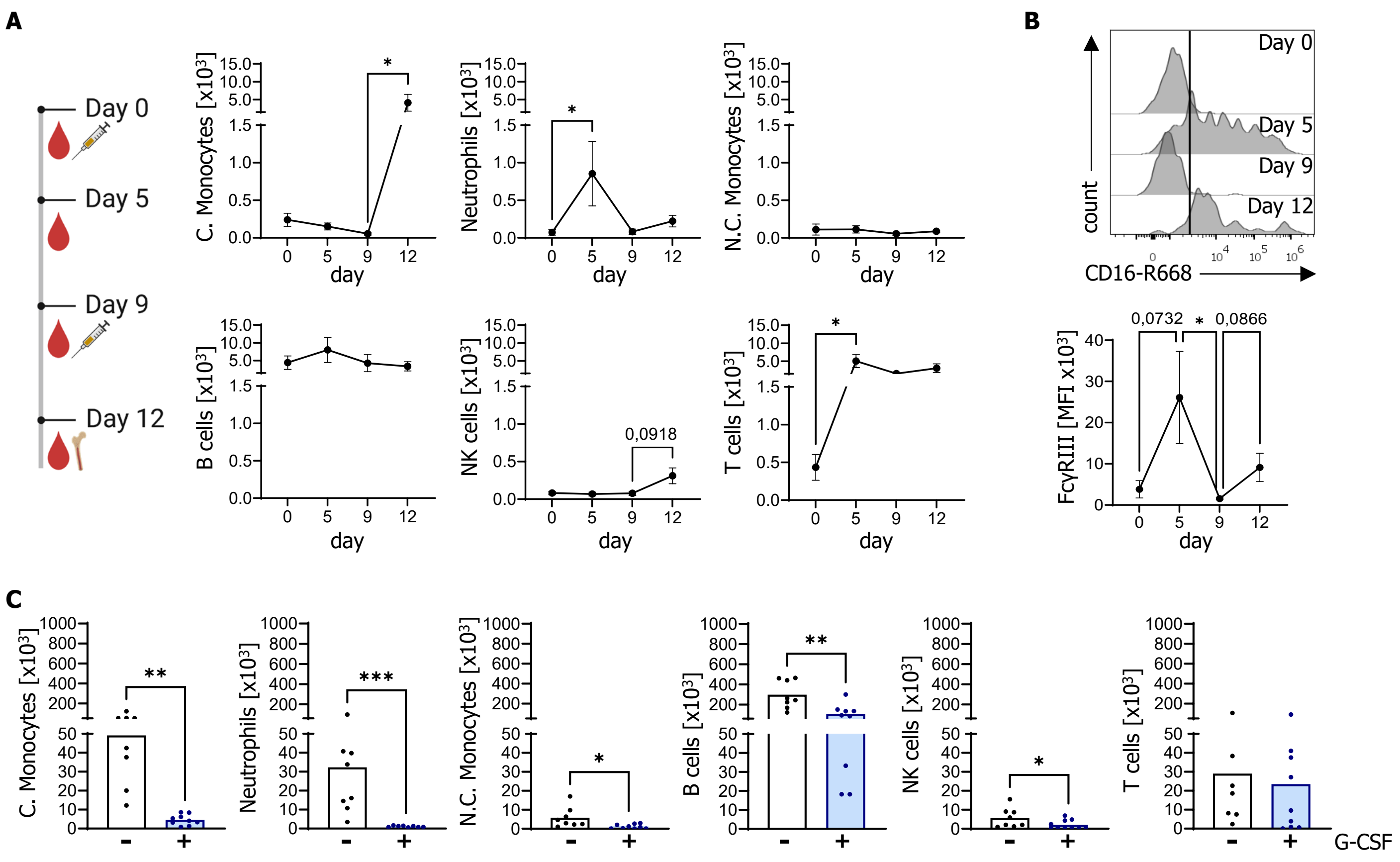


Fig. S4. **Human G-CSF is increasing and maturing human myeloid cells in the blood of HIS mice.**
(**A**) G-CSF was injected s.c. on day 0 and day 9. Human immune cell composition in the blood is shown (cells x$10^3$ per 100 µl blood). C., classical, N.C., non classical. (**B**) Quantification of human FcγRIII expression (CD16) and representative histograms. MFI, median fluorescence intensity. **(C)** Quantification of human immune cells in bone marrow samples by flow cytometry after G-CSF injection (cells x$10^3$ per femur). C., classical, N.C., non classical. Statistical analyses were performed using Shapiro-Wilk normality test. Depending on Gaussian distribution, samples were analysed using Ordinary one-way ANOVA or Kruskal-Wallis test, respectively (A,B). Depending on Gaussian distribution, samples were analysed using unpaired t-test or Mann-Whitney test (C). *$p < 0.05$; **$p < 0.01$; ***$p < 0.001$.

### Suppl. Figure 5

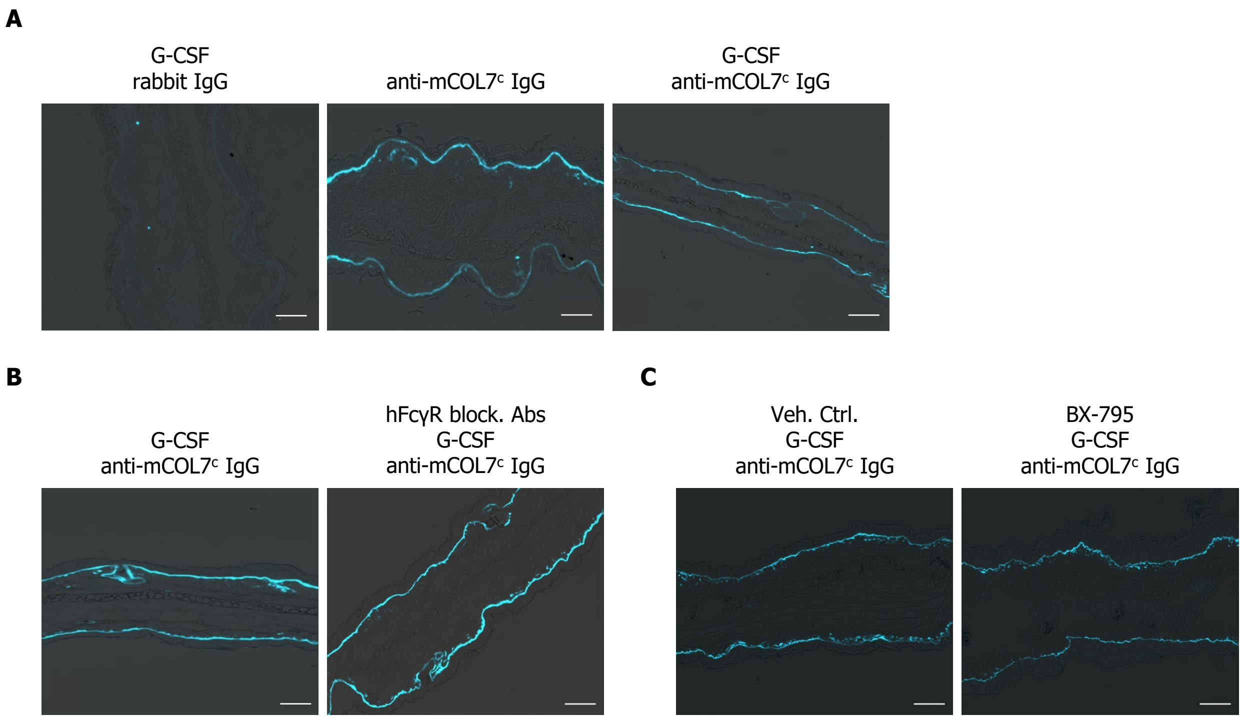


Fig. S5 **Deposition of rabbit anti-COL7$^{c}$ IgG in skin of HIS mice**
(**A**) Immunofluorescence staining of rabbit IgG deposited along the dermal-epidermal junction in ear samples of HIS mice injected either with total rabbit IgG (negative control), anti-mCOL7$^{c}$ IgG or anti-mCOL7$^{c}$ IgG and human G-CSF. (**B**) Immunofluorescence staining of rabbit IgG deposited along the dermal-epidermal junction in ear samples of HIS mice treated with human FcγR blocking antibodies. (**C**) Immunofluorescence staining of rabbit IgG deposited along the dermal-epidermal junction in ear samples of HIS mice treated with PDK1 inhibitor BX-795. The images were taken at 10× magnification, with the scale bar corresponding to 100 µm.

### Suppl. Figure 6

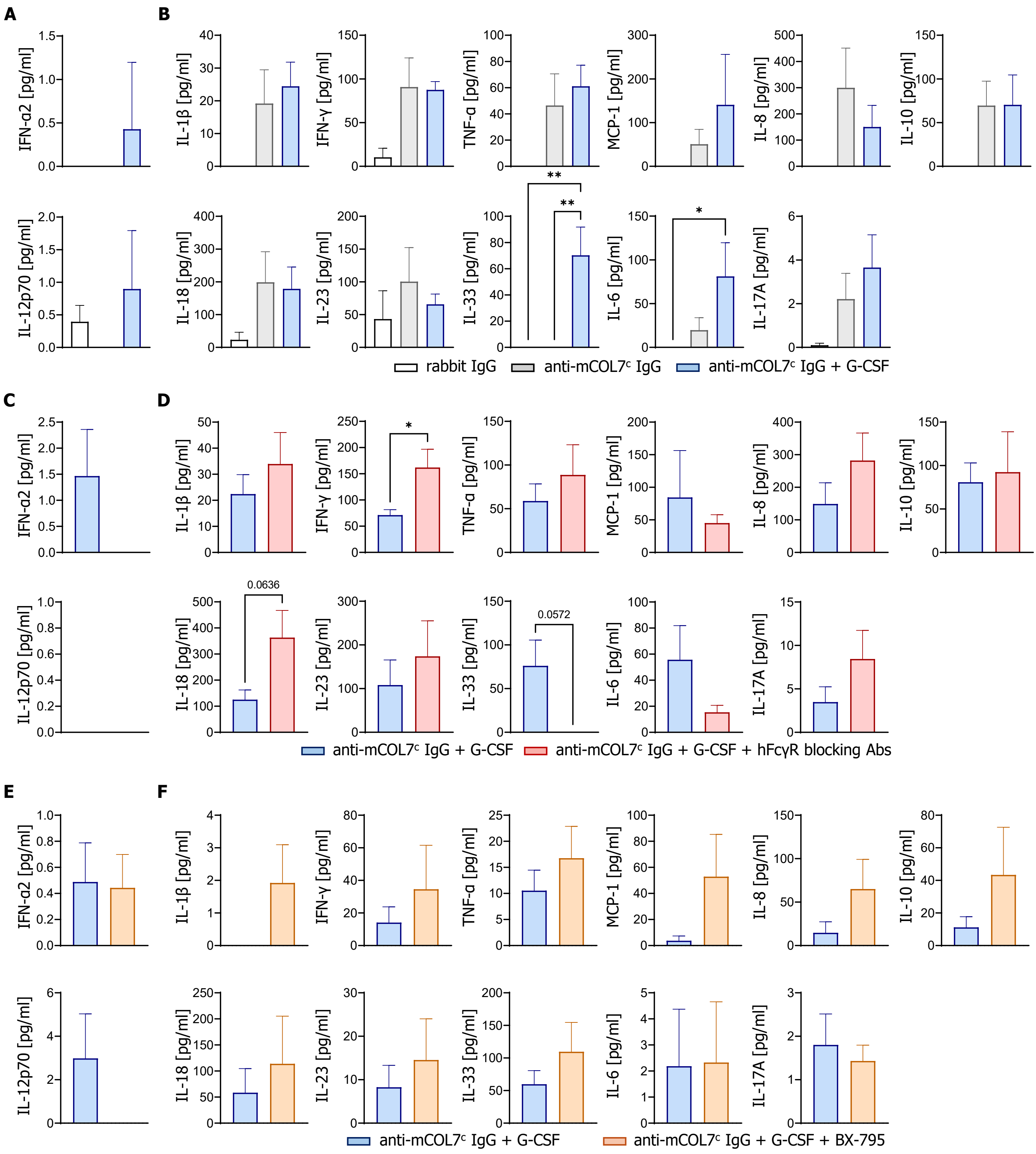


Fig. S6. **Human cytokines in the plasma of HIS mice after EBA induction.**
(**A**) Remaining human cytokins (day 13) in the plasma of rabbit IgG and anti-mCOL7ᶜ IgG treated HIS mice with and without human G-CSF injections. (**B**) Human cytokine levels in the plasma on day 6 of EBA HIS mice. (**C**) Remaining human cytokines (day 13) in the plasma of mice treated with and without blocked human FcγRs. (**D**) Human cytokine levels in the plasma on day 6 of EBA HIS mice. (**E**) Remaining human cytokines (day 6) in the plasma of EBA HIS mice treated with BX-795 or vehicle control. (**F**) Human cytokine levels in the plasma on day 13 of EBA HIS mice treated with BX-795 or vehicle control.

### Suppl. Figure 7

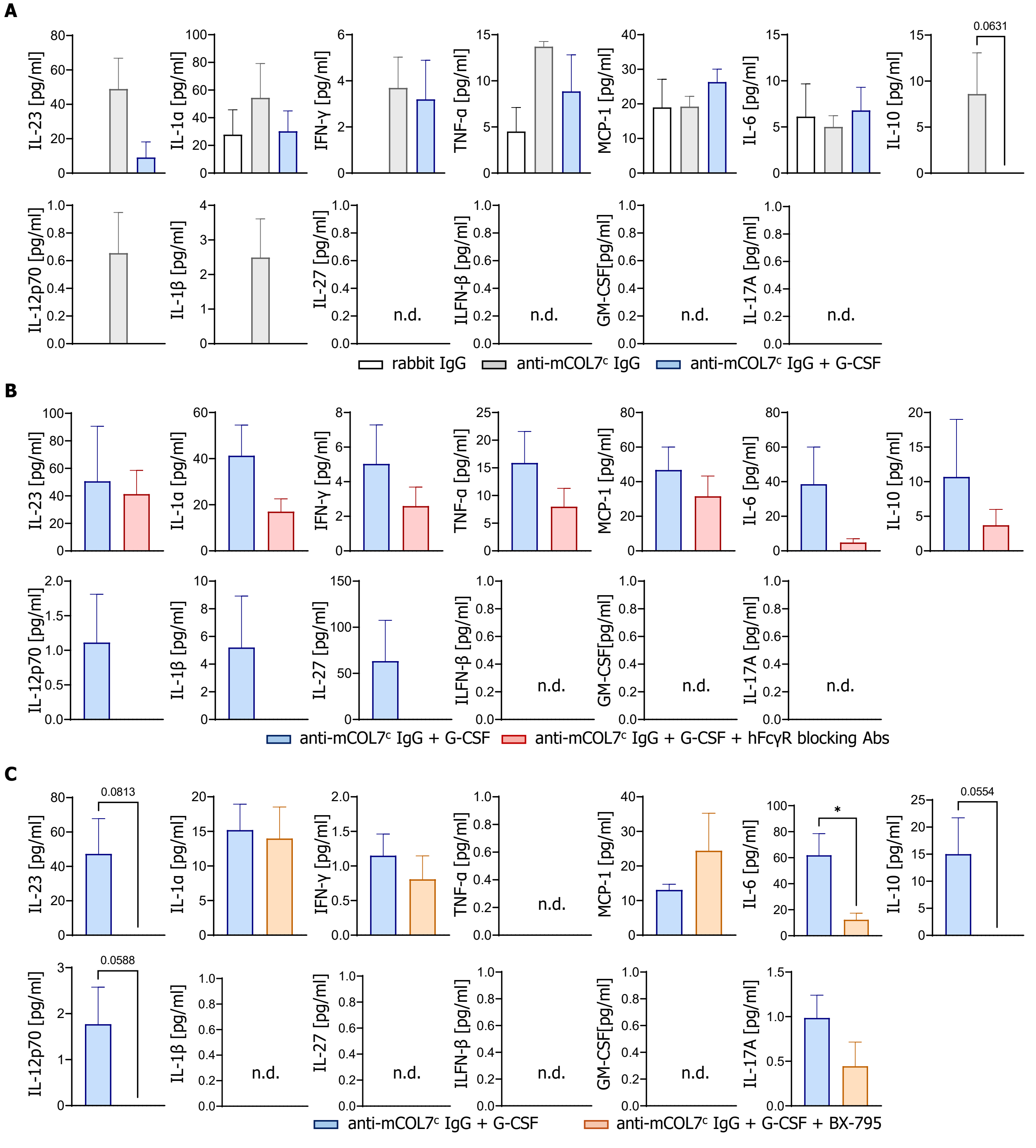


Fig. S7. **Murine cytokines in the plasma of HIS mice after EBA induction.**
(**A**) Murine cytokine levels in the plasma of rabbit IgG and anti-mCOL7c IgG treated HIS mice with and without human G-CSF injections. **(B)** Murine cytokine levels in the plasma of EBA HIS mice treated with and without blocked human FcγRs. **(C)** Murine cytokine levels in the plasma of EBA HIS mice treated with and without BX-795. n.d., non detectable.

### Suppl. Figure 8

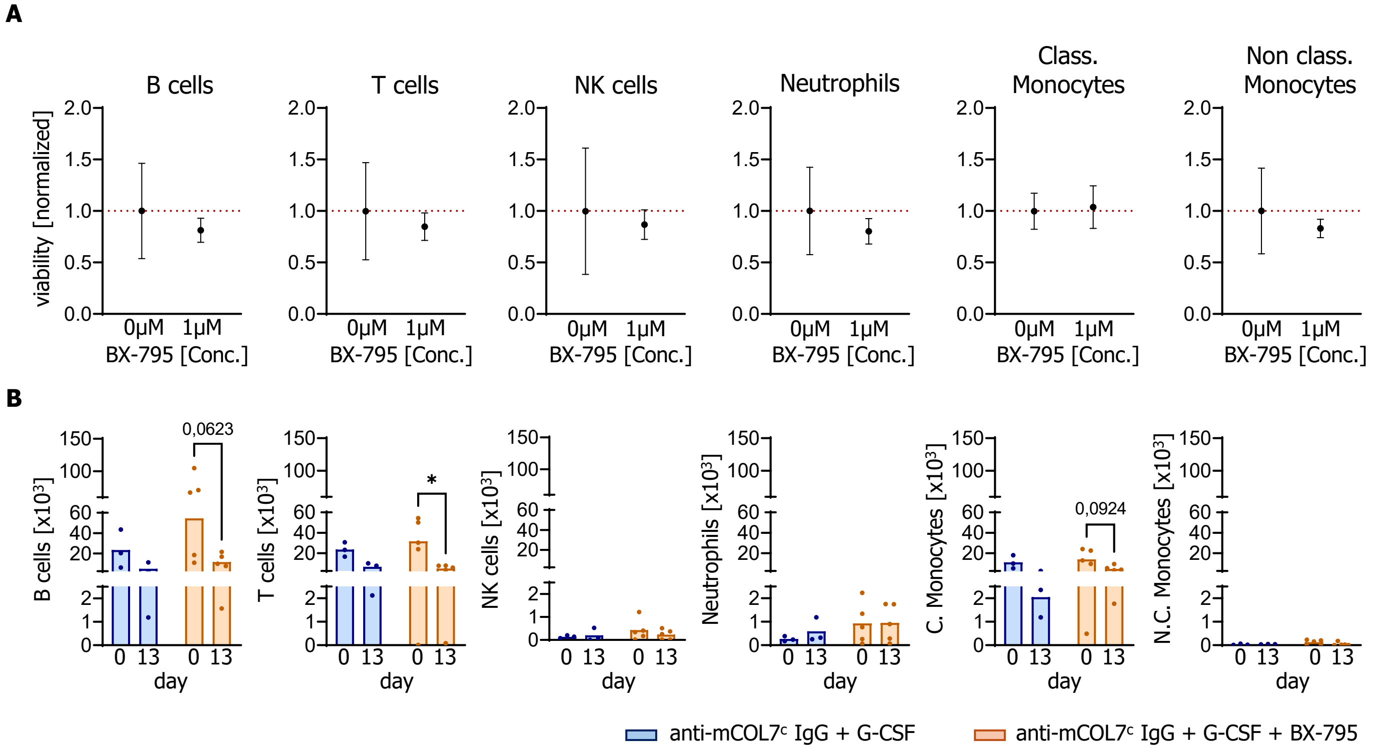


Fig. S8. **Viability of human cells treated with BX-795 in vitro and in vivo.**
(**A**) In vitro analysis of viable human cells treated for 30 minutes with 1 µM BX-795. Values are normalized to untreated controls. Data are shown as Mean ± SEM for 3 donors. Conc., concentration, Class., classical. (**B**) In vivo analysis of viable human cells in the blood of EBA HIS mice treated with BX-795 daily for 12 days. Cell numbers ($x10^3$ per 100 µl blood) of peripheral immune cells are compared from day 0 to day 13 for untreated and treated mice. In (A) Shapiro-Wilk normality test was used. Depending on Gaussian distribution, samples were analysed using paired t-test or Wilcoxon-test. Data from (B) was analyzed using Sidak's multiple comparisons test. *$p < 0.05$.
